# msCNVS: medium throughput single cell copy number variation sequencing with barcoded library construction free of preamplification toward clinical implementation

**DOI:** 10.1101/2024.04.01.587505

**Authors:** Guanchuan Lin, Bin Peng, Caiming Chen, Zhanying Dong, Mengchang Xu, Jinyu Gao, Jie Yu, Bei Jia, Chen Luo, Rui Hua, Changtai Xiao, Linlin Wang, Liyao Mai, Yulong Zhang, Yuanfang Lu, Yuanqiao He, Yali Song, Sadie L Marjani, Weimin Zhang, Junxiao Zhang, Mei Zhong, Song Quan, Sherman M Weissman, Hao Zhu, Xinghua Pan

**Author notes:** These authors contributed equally to this article (Guanchuan Lin, Bin Peng, Caiming Chen, Zhanying Dong). Corresponding authors: E-mail: Xinghua Pan, Hao Zhu.

## Abstract

Single cell copy number variation sequencing (scCNV-seq) is valuable for genomic analysis of a variety of health and disease systems, yet the available methods either depend on either preamplification of the whole genome of each cell, special devices or untracable, which hinder scCNV-seq practice in clinics. Here we provide scalable multiplex scCNV-seq (msCNVS) that allows direct medium-throughput library construction for single cells, which are barcoded individually with Tn5 transposome in a microplate and immediately pooled for downstream processes, elevating the efficiency by orders of magnitude. An algorithm is developed on the distribution of segmented normalized read count density to identify the major peaks associating with integer copy numbers, further improving the accuracy for objective CNV calling. Here msCNVS faithfully distinguishes the unique CNV patterns of 5 cell lines. The individual msCNVS profiles of 70 individual K562 cells highly correlate with the profile of K562 bulk cells (R=0.90-0.98), and the triplicates of HeLa3 bulk cells correlated nearly perfect (R=0.99). The msCNVS CNV profiles of primary cell cultures of amniotic fluid are confirmed by G-banding karyotype analysis and chromosomal microarray analysis. Additionally, the coverage uniformity of msCNVS is superior to that of MDA and MALBAC and approaches what eMDA and DOP-PCR achieves. Furthermore, msCNVS detects CNV deletions in 2 stunted, abnormal blastocysts, one with CNV mosaicism, and uncovered variants of CNVs in circulated tumor cells, cancerous pleural effusion cells, and patient-derived xenograft nuclei. Thus, msCNVS promises a robust, reliable and highly efficient approach to genomic testing of precious and rare cells like those typically obtained in reproductive and cancer clinics.

**One-Sentence Summary:** By early pooling of multiple single cells individually barcoded with Tn5 transposome, msCNVS enables efficient CNV profiling of rare clinic samples

## 1. Introduction

Copy number variation sequencing (CNV-seq), especially single cell CNV-seq (scCNV-seq), has revealed the landscapes of cancer evolution and embryo mosaicism in recent years; however, its clinical application for molecular diagnosis using embryo or cancer liquid biopsies lags behind^1–3^. Chromosomal or segmental aneuploidy (involved in different scales of CNVs, or called CNAs, i.e., copy number alternation, when initiated in somatic cells) is one of the most important factors attributed to early pregnancy failure and miscarriage^4^. Conventional prenatal diagnosis (PD) primarily utilizes chromosome microarray analysis (CMA) of amniotic fluid or chorionic villus cells and traditional chromosome karyotyping. Both non-invasive prenatal testing (NIPT), by sequencing cell-free DNA isolated from maternal plasma, and preimplantation genetic testing (PGT), by sequencing a biopsy composed of a few trophoblast cells, have been in practice for some time^5^. While assessment of single nucleotide variation (SNV) associated with Mendelian diseases and CNVs by CMA is reliable, more and more CNV-seq tests are applied but on bulk or micro-bulk cells. NIPT and PGT will be more accurate and efficient at detecting mosaicism if scCNV-seq is employed and analyzed robustly ^6,7^.

Similarly, a number of studies evaluating the CNV of circulating tumor cells (CTC) or minimal residual disease (MRD) have been reported but few in clinical practice, likely due to unsatisfactory robustness and efficiency in both single cell CNV detection and CTC isolation^8–10^. Compared with SNVs of mutation burden, which are well recognized in cancer precision medicine, CNVs may be another critical contributor leading to malignant progression for both solid cancers and some leukemia, and CNV identification may result in more effective biomarkers for early screening, monitoring and guiding precision cancer therapy^11,12^.

The initial efforts for scCNV-seq were based on the preamplification of the whole genome (preWGA) ^1^, either in complete or sparse coverage, of individual cells, followed by library construction of each amplicon and next-generation sequencing (NGS). These efforts include degenerate oligonucleotide-primed PCR (DOP-PCR)^13^, multiple displacement amplification (MDA)^14^, multiple annealing and looping-based amplification cycles (MALBAC)^15^, and linear amplification via transposon insertion (LIANTI)^16^. These preWGA-based scCNV-seq methods enable the processing of the genome of single cells, after WGA, and have been applied widely for scCNV-seq with many fruitful results^17^. Some WGA methods are less biased and give faithful CNV calling, including examples eMDA^18^, MIDAS^19^, LIANTI^16^. However, with the preWGA amplicons, the downstream library construction is low throughput, resulting in low labor efficiency, high time consumption, and high cost. Critically, preWGA derived biases are often introduced in data, and they cannot be removed using computational methods ^20^.

Since Adey et al. (2010) reported a modified Tn5 transposon^21,22^, a versatile approach enabling sequencing library construction directly from single cells without preWGA^20^, single cell sequencing capability has greatly accelerated. Adey and colleagues developed single cell combinatorial indexed sequencing (SCI-seq)^23^, followed by other derivatives, such as sci-L3^24^ and sci (‘s3’)^25^. However, with these high-throughput methods, the original identity of each single cell, which is important for rare and precious cells, is missing from the output data. Meanwhile, Zahn et al. (2017) presented direct library preparation (DLP) using Tn5 transposome in nanoliter-volumes for scCNV-seq^20^, which also skips preWGA and improves uniformity and efficiency. However, DLP builds libraries separately for each cell and depends upon a special microfluidic device or droplet emulsification. In addition, scRNA-seq provides another high-throughput way to reveal cancer architecture by inferring CNVs in single cells^26^. These approaches have led to exciting discoveries by answering foundational biological questions, such as the initiation sites and dynamics of DNA duplication and somatic CNV patterns in variants of cell lineages during aging^20^. They also promote clinical applications, particularly in how aneuploidy profiling of an embryo is associated with pregnancy outcome, and how CNV patterns of CTCs determine cancer progression^2,27,28^. These studies provide valuable foundation and lessons in data analysis and technique setting for current exploration.

In research of genetic and somatic diseases, cell number is usually large and high throughput is preferred. However, in clinical analysis of liquid biopsy from cancer, or biopsy from an embryo or fetus, the cells are often rare and precious^29,30^. Conventionally, the CNA profile analysis for a single cell isolated from CTCs or a few cells from a preimplantation embryo relies on preWGA. Although numerous improvements have been made for preWGA-based methods, they are still plagued by low efficiency and issues with bias^30^. The ideal scCNV-seq method should be precise and reliable at the industrially required scale, efficient in time and cost, robust and friendly in operation, and capable of tracing the identity of each cell input from the sequencing output^31,32^.

We have developed an approach that fulfills the above requirements particularly for clinical practice. This method, termed msCNVS, uses conventional microplates or strip tubes, to reach a medium-throughput and is scalable to high-throughput when adapted to a microfluidic or droplet system. Based on early barcoding of each single cell genome with a modified transposome and immediate pooling of multiple single cells, msCNVS empowers single-tube library construction with multiple single cells. The output library is compatible with current NGS platforms, particularly Illumina, without any adjustment of procedure or reagents. Combined with a renovated algorithm that transforms the frequencies of segmented-normalized-read-counts to a density distribution to obtain a more objective CNV calling that is based on the distribution peaks associated to the integer ploidy levels, the reliability and feasibility of msCNVS are demonstrated with a panel of HapMap cell lines. We further verified the technique on single cells from amniotic fluid cell culture and compared the results to conventional G-banding karyotype and CMA analyses. msCNVS was also employed on single cells isolated from abandoned blastocysts, and marked heterogeneity of chromosome aneuploidy or mosaicism was identified. Finally, msCNVS was used to analyze CTCs enriched from peripheral blood of colorectal cancer (CRC) patients and pleural effusion cells from non-small cell lung cancer (NSCLC) patients, as well as patient-derived tumor xenograft (PDX) nuclei of CRC. In these important cancer applications, apparent CNAs and heterogeneity were revealed. Taken together, msCNVS provides a practical approach for genomic testing of diseases at single cell resolution, enabling the analysis of cells from embryos and fetuses as well as adults for germline and somatic CNV profiling.

## 2. RESULTS

### 2.1. msCNVS implements direct barcoding of individual genomes and early pooling

To develop an efficient and faithful assay for measuring CNVs simultaneously for a panel of single cells, we built an approach that involves tagmentation of each single cell genome using Tn5 transposome with a set of newly designed oligonucleotides. These oligonucleotides incorporate sample barcodes allowing early pooling of multiple single cells, while precisely tracing the identity of each cell. A pair of handles are used to enable efficient addition of the library indexes and tag sequences for amplification and priming during sequencing on the Illumina platform. The handles are asymmetrical in the 2 ME sequences of Tn5 transposome, and the 8-nucleotide barcodes rely on one of the 2 ME sequences. After lysis of cells in separate wells (each contain a single cell), Tn5 transposome is added for sequence tagmentation and cell barcoding. Next, the contents of all wells are pooled into a single tube. Then, a common PCR program captures the handle sequences flanking each fragment from each cell for all cells in a pool for sequencing.

The workflow of msCNVS is briefly shown in Fig. 1a. To summarize, a single cell suspension is prepared, and the desired target cells are delivered into an 8- or 12-well strip or a 96- or 384-well plate, by a manual or robotic pipette or flow cytometer. We currently perform 8-cells or 48 cells with barcodes×2 indexes (i5, i7) in a 96-well plate. After cells are thoroughly lysed in each well using Qiagen protease, the protease is deactivated, and the Tn5 transposome assembled with double-strand oligonucleotides is added to each well. In this step, the whole genome of each cell is evenly fragmented, and each fragment in a pool of fragments is flanked by ME sequences-containing barcoded oligonucleotides. The panel of cells are then pooled in a single tube for PCR amplification to complete the final library construction step with a pool-specific index built into the PCR primer. Finally, the libraries are sequenced, independently or in combination with other libraries on a dual-index sequencing platform, which in this report is Illumina NovaSeq. The required output raw sequencing data is typically 1-3Gb per cell. In contrast, the raw data for REPLIg or MALBAC-based scCNVS is usually much larger; for example, the available data for a human umbilical vein endothelial (Huvec) cell ranges between 30-50 Gb^18^.

**Fig. 1.**
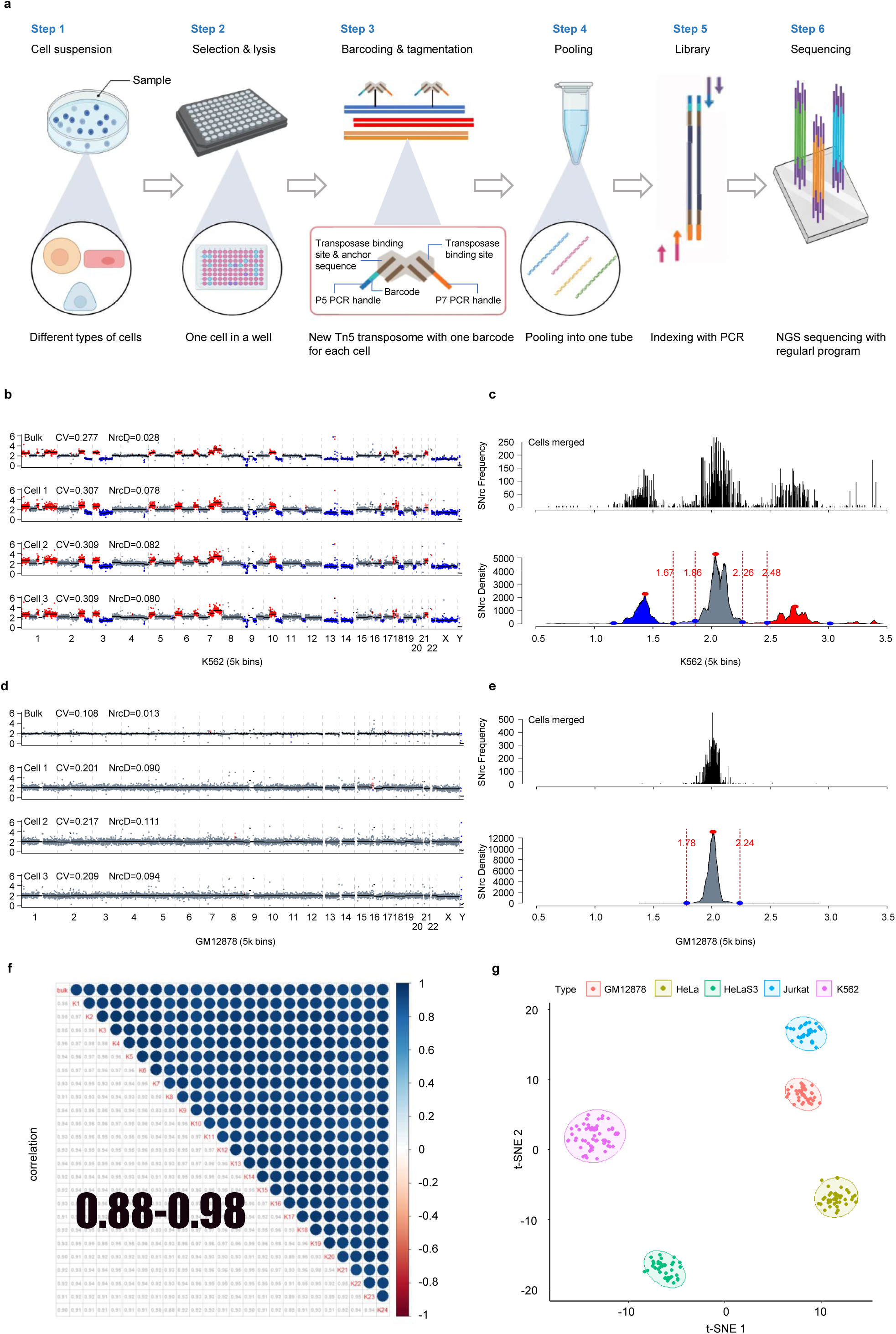
CNV calling of single cells by mcCNVS is specific and consistent to bulk. **a. Workflow of msCNVS.** Single cell suspension generated. Individual single cells are delivered into each well of a 96-well PCR plate or a strip tube. After cell lysis, a specially designed Tn5 transposome with P5Tn5 adapter and P7Tn5 adapter is added to each well for tagmentation and barcoding of all fragments over the genome of each cell. The DNA fragments of the panel of cells are then pooled in a single tube. PCR is performed for integration of the indexes and sequencing elements for the pool of fragments, which are subject to NGS. **b, c, d, e.** the X-axis represents the cross-genome copy number distribution. Y-axis represents the SNrc frequency of the given copy number over the whole genome. Red dot is the top of the major peak for each integer copy value. Vertical red dot line gives the boundaries of each major peak region. Blue: 1-copy or less; Purple: fuzzy deletion (value between 1-copy and 2-copy); Grey: 2-copy; Pink: fuzzy duplication (value between 2-copy and 3-copy); Red: 3-copy or more. **b. CNV patterns of K562 cells.** The calling strategy is described in the Methods. The first row is K562 bulk from ENCODE database, and the second to fourth rows are K562 single cells by msCNVS. **c. SNrc frequency distribution and SNrc density distribution of the merged data for K562 single cells. d. CNV patterns of GM12878 cells.** The first row is GM12878 bulk from American National Library of Medicine database, and the second to fourth rows are GM12878 single cells by msCNVS. **e. SNrc frequency distribution and SNrc density distribution of the merged data for GM12878 single cells. f. correlation between single cells and bulk data of K562.** The CNV patterns of a set of 24 selected K562 single cells by msCNVS are highly correlated to each other (R=0.88-0.98), and to the corresponding bulk (R=0.90-0.95). The resolution is 5k bins/genome, or 0.66 Mb/bin. **g. tSNE clustering for the five cell lines based on the msCNVS patterns.** Each of the 5 cell lines assayed by msCNVS, GM12878, K562, Jurkat, HeLa and HeLa S3, grouped into a distinguished subcluster through tSNE according to their CNV patterns. Two isologous cell lines, HeLa and HeLa S3, are also distinct from each other by their CNV profiles.

This approach, termed msCNVS (multiple single cell CNV-seq), is exceptional in several ways. It avoids preWGA for each cell, which is usually applied in a variety of methods before library construction. A unique sample barcode is incorporated in each cell for early pooling, increasing the efficiency and specificity of library construction and enabling the identity deciphering of the cells after sequencing. Because of the basically random fragmentation caused by direct Tn5 tagmentation, each fragment is unique in the terminal sequences, such that any overlapping capture of a given DNA segment are recognized, and duplicate library products are removed computationally from the raw sequencing data (Extended Data Fig. 1b), which is impossible with the preWGA approaches. These features not only reduce the cost, but also improve the faithfulness of CNV calling. Taken together, msCNVS implements a scalable-throughput analysis of single cells and generates representative libraries of their original genomic templates, enabling parallel and accurate calling of CNVs and trace to the origin of the cells (Extended Data Fig. 1a).

### 2.2. Faithfully reflecting CNVs of single cells by msCNVS

We assessed the reliability of the CNV calling by msCNVS using both published references and our own bulk sample results. For CNV callings, we developed a Python pipeline based on the concept of variable-sized genomic windows ^33,34^. A minimum of 35 unique, high quality aligned reads per bin (i.e., median 35 reads/bin) were set, with the default bin number of 5k (or resolution of 0.66Mb) over the genome. We tested several cell types and clinical samples to evaluate and validate msCNVS. We first applied msCNVS to the human erythroleukemic cell line K562 and the lymphoblastoid cell line GM12878, both of which have been extensively examined for their CNV landscapes. With raw data 1-3Gb/cell (and final mean depth of ~0.03-0.06×genome, covering ~2-6% of the genome), the msCNVS profiles of both cell lines showed patterns that consistently matched the corresponding bulk results (Fig. 1b, d). In the K562 results, events of amplification in chromosomes 1, 2, 3, 4, 5, 6, 7, 10, 18, and 21, and deletions in chromosomes 2, 3, 4, 9,10, 11, 12, 13, 14,17, 18, 19 and 20 were observed; in these chromosomes the CNV sizes ranged from 1,177 Kb to 83,430 Kb. In the GM12878 results, no significant copy amplification or deletion was observed under the same analysis parameters (Fig. 1d).

It is desirable to robustly obtain a reliable and precise calling at a relatively low sequencing depth for cost efficiency. The aggregated msCNVS profile of 70 single K562 cells (24 respective single cells are showed in Fig 1e) was highly correlated with the bulk cell data retrieved from the ENCODE project (ENCSR025GPQ^35^) (R = 0.75 to 0.98, median =0.91) (Supplementary Table 1and Extended Data Fig. 2a). The msCNVS results of K562 single cells were also highly correlated with those from a micro-bulk analysis (100 to 1000 intact K562 cells) and purified K562 DNA at 10 ng and 50 ng, respectively, by mCNVS (Extended Data Fig. 2b, c). To access the influence of the heterogeneity among single cells, we further analyzed, in triplicate, of 500 HeLa cells using msCNVS; the correlation among them was 0.99 (Extended Data Fig. 2d, f), showing convincing reproducibility.

With a pseudo-bulk sample (silico-merged 70 single K562 cells), we got a total of 76Gb raw reads, which included 24,629,032 clean unique reads or at 0.897× genome depth, covered 38.83% of the genome, and qualified for CNV calling at the resolution of 5 Kb or 5k bins (Extended Data Fig. 4c and Supplementary Table 2). This pseudo-bulk analysis greatly raised the resolution over individual single cells (Extended Data Fig. 4), and allowed for the accurate assessment of the CNV breakpoints for a panel of single cells merged within the same cluster.

In addition to K562 and GM12878, we also analyzed single cell CNV profiles of cell lines HeLa, HeLa S3, and Jurkat using msCNVS (Extended Data Fig. 5c). The rate of successful library construction for cells from each cell line was 96%-100% (ex. HeLa, 48/48; HeLa S3, 47/48; Jurkat, 32/32). PCA (Extended Data Fig. 5a) and tSNE clustering (Fig. 1g) of the msCNVS results for the 5 cell lines gave an outstanding separation of five distinct clusters, each including all individual cells of their belonged cell line. The top 5 marker CNV areas marked the cluster identities (using the *Seurat* package) (Extended Data Fig. 5b). It is notable that when msCNVS was applied to HeLa and HeLa S3 cells, two very closed cell lines, the clusters were obviously distinct (Fig. 1g).

### 2.3. Compelling uniformity of normalized read counts (Nrc) in msCNVS

We evaluated the uniformity of different CNV approaches for the diploid cell lines Huvec and HT-29. Taking HT-29 as an example (Extended Data Fig. 6), the correlation between 8 single HT-29 cells and its bulk counterpart obtained by msCNVS and mCNVS, respectively, was 0.8. In this case, both were better than MDA, with correlation of 0.64 and 0.71, and MALBAC, with correlation of 0.68 and 0.74^18^. We noted that the HT-29 bulk sequencing data we generated was slightly different in chromosome 1 from the bulk data of a database^18^, which could be attributed to some new mutations in the HT-29 during passaging in culture. For HT-29 msCNVS results, the deviation values, CV and NrcD, were slightly higher than that generated by DOP-PCR, and significantly lower than that by MALBAC and MDA (Extended Data Fig. 6a). When using Huvec cells as an example, the CV and NrcD for msCNVS were equivalent to that by eMDA, but apparently lower than that by MALBAC and MDA (Fig. 2a).

**Fig. 2.**
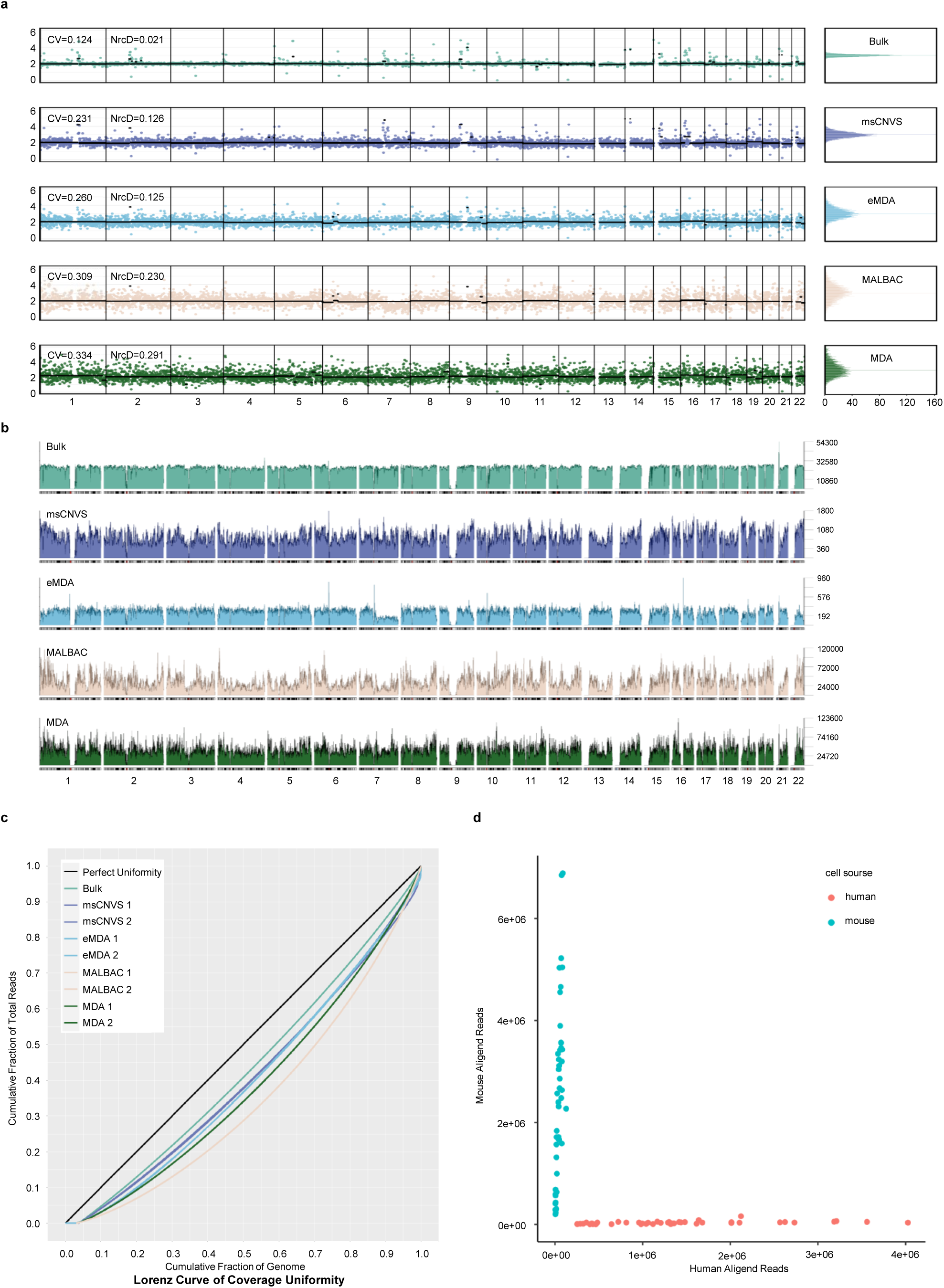
Coverage and uniformity of msCNVS in comparison with other CNV-seq methods. **a, b, c.** Single cell CNV patterns of Huvec by msCNVS, compared to those by eMDA, MALBAC and MDA (with reference bulk data) extracted from publicly available sources. The genome is divided into 5k bins (0.66 Mb/bin) in the CNV calling. **a. comparison of deviations for CNV calling displayed with different methods.** Here no CNV is displayed, consistent with the nature of the diploid cell line. Both CV and NrcD are listed for derivation measurement. **b. Genome-wide coverage and fluctuation.** The value of aligned reads in each bin, without normalization, is displayed directly along the genome. The sequencing depths are proportional to the Y-axis scales of the methods listed. **c. Lorenz curve of coverage uniformity.** Two single cells with the best data reported are showed for each method. **d. Clarification of cross-contamination among individual samples /cells.** Human and mouse single cells were individually barcoded and pooled in different batches of experiment (in total: 48 mouse C2C12 cells, and 44 human Huvec cells) for msCNVS library construction and sequencing. Subsequently, unique sequences were acquired and mapped to human and mouse genome in parallel. As a result, there is no evident cross-contamination of reads detectable.

msCNVS was also assessed with the Lorenz curve of coverage uniformity. For each of the 2 cell lines in the comparison, the best performing 1 or 2 cells from eMDA, MDA and MALBAC were selected from the NCBI Sequence Read Archive database (accession no. SRP052908). Similarly 1 or 2 cells after msCNVS with the raw read number similar to eMDA were chosen (Fig. 2b). Compared with scCNV-seq by MDA and MALBAC, msCNVS outperformed these methods in terms of signal uniformity or fluctuation. Specifically, the uniformity was closer to the bulk analysis with perfect curves for Huvec and HT-29 cells. msCNVS also outperformed MDA and MALBAC for HT-29 cells, although DOP-PCR performed slightly better than msCNVS(Fig. 2b and Extended Data Fig. 6b).

To examine any possible cell-barcoding cross-contamination in our CNV sequencing library that may average the signals among different cells in a pool and distort CNV calling, we used 48 (48/48 successful libraries) cells of the mouse cell line C2C12, and 44 (44/48 successful libraries) cells of the human cell line, Huvec, for pooling library construction in multiple batches. We found that all 92 mouse and human cells matched their corresponding reference genome (hg19 or mm10) well (Fig. 2d). No evident cross-contamination was observed.

In the msCNVS results of K562, HeLa, and HeLa 3S, we found obvious heterogeneity in these cellular populations (Extended Data Fig. 7). In line with this, heterogeneity of the K562 cell line from protein markers to RNA profile was previously reported ^36^. Furthermore, heterogeneity of HeLa^37^ and HeLa S3^38^ was revealed by scRNA-seq.

### 2.4. Sensitive detection of chromosomal or segmental aneuploidy with single cells from amniotic fluid cell cultures and preimplantation blastocysts

CNV analysis is a basic technique for screening aneuploidy after in vitro fertilization or during prenatal development. To test the ability of msCNVS in this field, we used msCNVS to assess single cells obtained from the primary cell culture of amniotic fluid from 5 fetuses with apparent CNVs (Fig 3 and Extended Data Fig. 8 a, b) and 1 healthy control (Extended Data Fig. 8 c). msCNVS identified three fetuses (Y9000 in Fig. 3d, g, Y9004 and Y9043 in Extended Data Fig. 8a, b) with trisomy 21, one fetus (Y9005) with segmental amplification on Chr 2 (Fig 3e, i), and a fetus (Y9007) with a deletion on Chr 18 (Fig. 3f, k). All of these CNVs were confirmed by G-banding karyotyping (Fig 3a, b, c) and/or chromosome microarray (CMA, Fig 3h, j, l). Importantly, the single cell CNV patterns revealed by msCNVS were generally homogeneous within each fetus (Fig. 3d, e, f). These results show that msCNVS obtained 100% success rate with all 8 single cells/per sample for the 3 samples with trisomy 21 (Fig. 3a, d, g, h and Extended Data Fig. 8 a, b). In addition, a chromosome abnormality (Chr 21 of fetus Y9043 was linked to Chr 16) was also detected precisely (Extended Data Fig. 8 b).

**Fig. 3.**
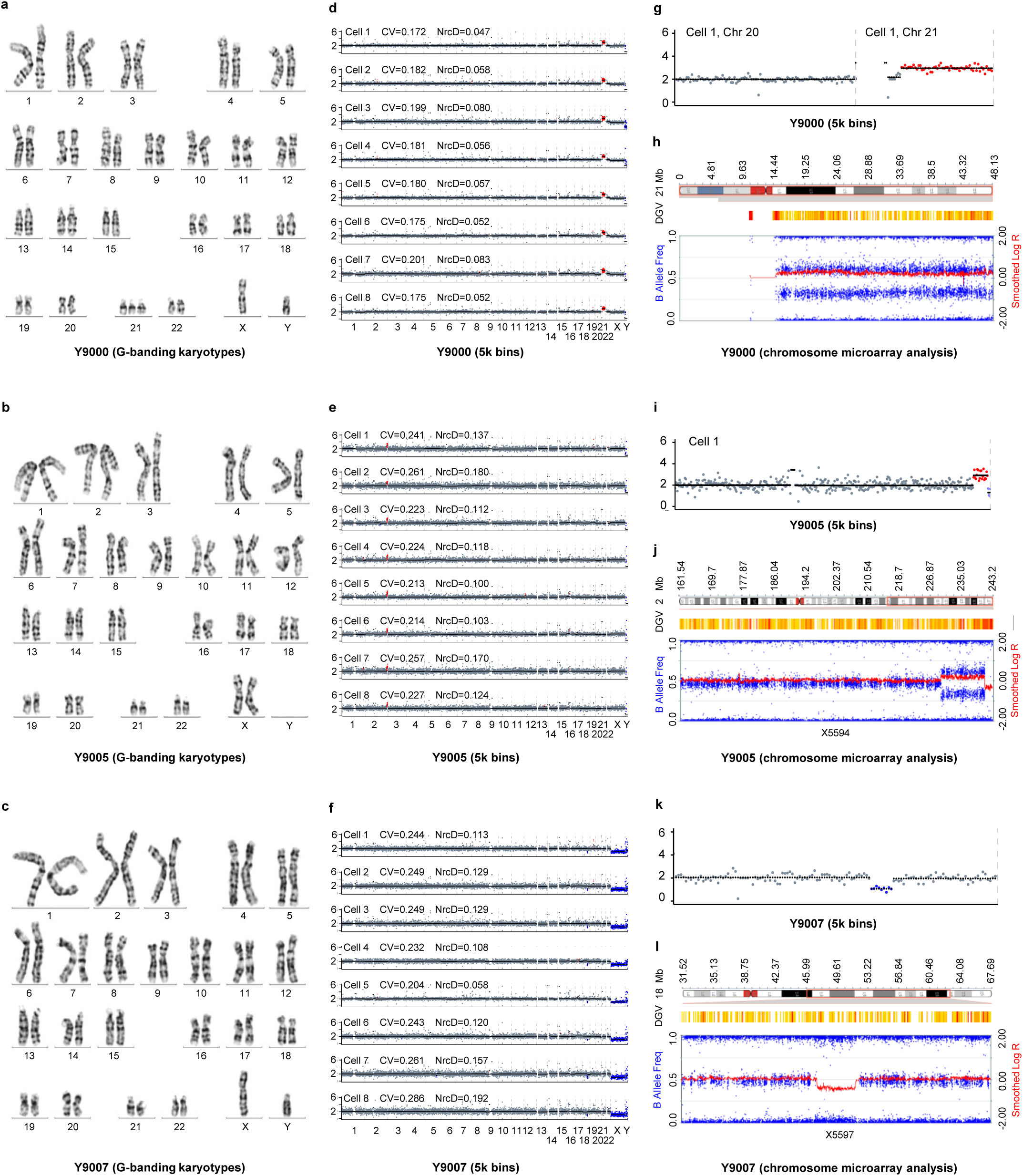
Assessment of msCNVS with single cells from primary amniotic fluid cell cultures of three fetuses. A resolution of 5k bins applies for msCNVS pattern. For eight cells from each of the three fetuses, all cells exhibited the CNVs on the corresponding chromosome **(d, e, f),** as the aneuploidy detected by the G-banding karyotypes (**a, b, c**). The detected CNVs are further highlighted (**g, i, k**) and validated by chromosome microarray analysis (CMA) (**h,j,l**). **a, d, g, h. fetus Y9000 with trisomy 21 (3-copy)**. **b, e, i, j. fetus Y9005 with a segmental amplification and a segmental deletion on Chr 2 long arm**. i.e., a duplicated segment (3-copy) in Chr 2: 229,613,850 to 240,229,580 is unveiled by msCNVS, with a missing copy (1-copy) in Chr 2: 240,818,881 to 242,593,602. These are confirmed in the CMA detecting a segment duplication (3-copy) in Chr 2: 2q363q37.3 (Chr 2: 229,565,839-241,021,460) duplication, and a segment missing (1-copy) on Chr 2: 2q37.3 (Chr 2: 241,023,450 - 243,044,147). **c, f, k, l. fetus Y9007 with a segmental deletion (1-copy) on Chr 18 long arm**, i.e. Chr 18: 47,292,199 to 54,971,061 is detected by msCNVS. Consistent with it, CMA shows one copy missing on Chr 18: 18q21.1q21.2 (Chr 18: 47,244,725-51,866,974).

In fetus Y9005, CMA showed that Chr 2 at 2q36.3q37.3 was duplicated for one additional copy, and the size of the duplicate fragment was about 11.45 Mb. Meanwhile, one copy of a region of Chr 2 at 2q37.3 was missing, and the size of the missing fragment was about 2.02 Mb. Similarly, msCNVS detected a duplicated copy of the segment between 229,613,850 to 240,229,580 of Chr 2, and one missing segment between 240,818,881 to 242,593,602 of Chr 2 (Fig. 3i, j). In fetus Y9007, CMA showed that one copy of Chr 18 at 18q21.1q21.2 was missing, and the size of the missing fragment was about 4.52 Mb. msCNVS not only identified the same Chr 18 result, but also precisely indicated the missing genes, including *CDCC11*, *DCC*, *MYO5B* and *SMAD4* (Fig 3k, l). As a control, eight cells with normal karyotypes were assessed with msCNVS. They were all diploid (Extended Data Fig. 8c). Therefore, all these CNVs, including duplication or/and deletion on Chr 21, Chr 18 and Chr 2, were identified by msCNVS and confirmed by G-banding karyotyping and/or CMA.

Next, we applied msCNVS to a set of single cells isolated from 2 blastocysts discarded due to poor quality (Fig. 4 and Extended Data Fig. 9). Blastocyst 2 was shown to contain the distinct cell subpopulations that likely originated from a common precursor and later diverged phylogenetically. The obvious mosaicism as identified in Blastocyst 2 was recently reported ^2,39^, yet the mechanism and the clinical significance need further investigation.

**Fig. 4.**
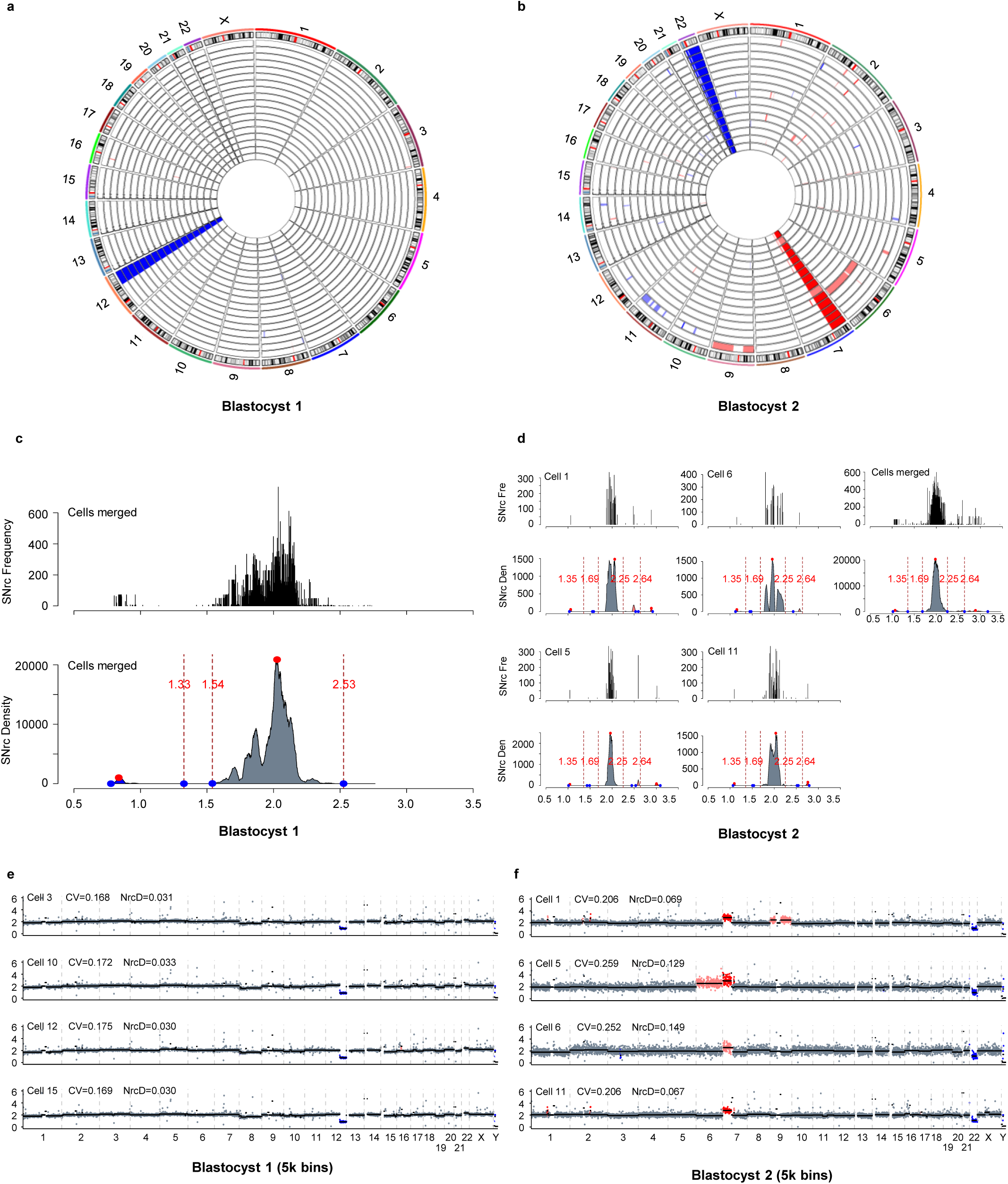
msCNVS patterns of single cells from two discarded blastocysts due to poor quality. The color labels are the same as in Fig. 1. A resolution of 5k bins applies for msCNVS pattern. **a, b, Circle map of the CNV profiles of blastocyst 1 (a) and 2 (b).** Each ring represents the chromosomal CNV profile of a cell. **c, d. SNrc frequency distribution and SNrc density distribution of single cells from blastocyst 1 (c) and 2(d). e. The CNV patterns of 4 representative single cells of blastocyst 1.** Including these cells, all 18 cells (Extended data Fig. 9a) are homogeneous, with a copy missing on the short arm of Chr 12 (1-copy). **f. The CNV pattern of 4 representative single cells of blastocyst 2.** Including these cells, all 15 cells exhibit segmental trisomy on the short arm of Chr 7, and disomy on the entire Chr 22 (Extended data Fig. 9b), while one cell, blastocyst 2 cell 1, exhibits additional segmental trisomy on Chr 9, and another cell blastocyst 2 cell 5 exhibits additional segmental trisomy on Chr 6.

Overall, msCNVS provides more accurate and highly efficient detection of abnormal chromosomes in embryos and fetuses, which can hopefully be applied to clinical single cell preimplantation genetic testing (PGT) and prenatal diagnosis (PD).

### 2.5. CNV assessment of single, circulated tumor cells (CTCs) from peripheral blood and cancerous pleural effusion cells, and nuclei from PDX tissue

To determine the potential of msCNVS in the analysis of liquid biopsies of cancer, we utilized the method to analyze the CTCs isolated from cancerous pleural effusion cells of 2 patients with non-small cell lung cancer (NSCLC 1, NSCLC 2) and CTCs from the peripheral blood of a patient with colorectal cancer (CRC 1) (Fig. 5a, b, c, d, e and Extended data Fig. a, b, c, d). The CTCs were enriched and identified with immune-capturing and digital detection (Fig. 5a). It is notable that the CNV patterns of different CTCs from each patient typically shared a patient-specific, common CNA pattern, as only minor variations (particularly some segmental fuzzy zones) were demonstrated among the CTCs from the same patient. This suggests that these cells might be derived from a common progenitor. However, the CNA patterns of different patients that we analyzed were different overall, even between the 2 NSCLS patients. This reflects some heterogeneity between different NSCLC patients, which may be ascribed to different tumor stages, classification, or treatments. The pattern and mechanism underpinning the dynamics and heterogeneity of CTCs along the longitudinal course of a given patient, as well as in different patients, are worth systematic study with large cohorts, considering the potential in guiding precision cancer therapy ^32^.

**Fig. 5.**
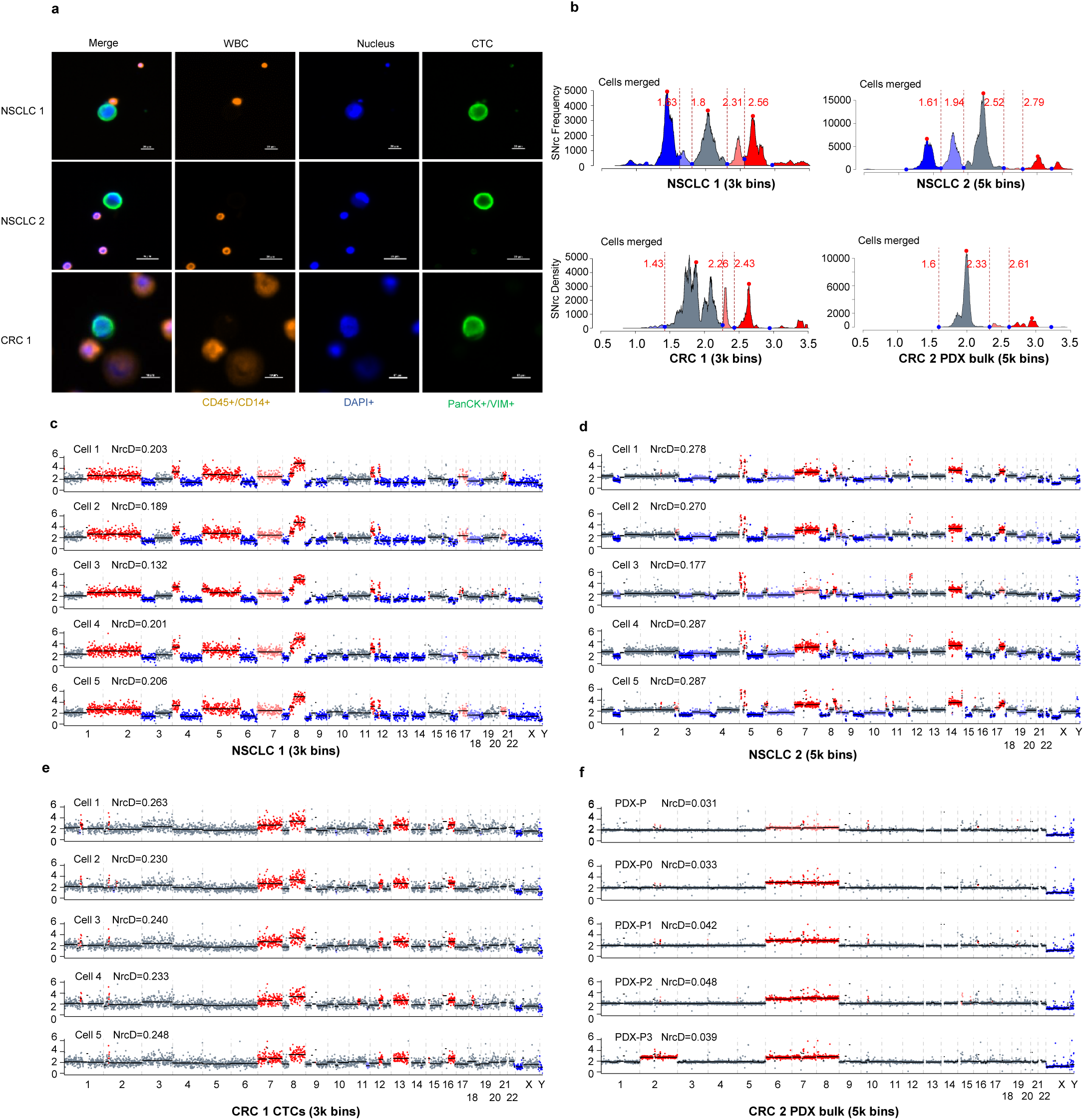
msCNVS patterns of single CTCs and bulk cells from NSCLC and/or CRC. The color labels are the same as in Fig. 1. 5k bins resolution were applied in (d, f), while 3k bins applied in (c,e) to reduce the derivation of SNrc. **a, Immuno-staining validation of CTCs** captured from human NSCLC and CRC on microfluidic HB CTC-Chip, versus normal leukocyte (WBC). CTCs are positive staining for cytokeratin PanCK or VIM or both (all green), without CD45 and CD14 staining (orange). WBCs are those cells with positive CD45 or CD14 staining (orange/PE). The nuclear of both CTCs and WBCs are DAPI staining (blue). **b, The SNrc density distribution** based on SNrc frequencies merged all samples analyzed in each of the 4 patients, NSCLC 1 (c, and Extended data Fig 10a, b), NSCLC 2(d and Extended data Fig 10c), CRC 1(e and Extended data Fig 10 d, e), and CRC 2 (f). **c, d, e: The CNV patterns** for single CTCs. Five single CTCs from each patient, NLCLC 1 in 3k bins (c), NLCLC 2 in 5k bins(d) and CRC 1 in 3k bins(e) are displayed. The CNV patterns are different from one to another patient. All single CTCs from the same patient share a common pattern for most CNAs, while some CNAs vary somewhat from cell to cell especially those in segmental fuzzy zones. **f. The CNV patterns for 5 bulks of PDX.** Each obtained from one generation of PDX tissue derived from CRC 2 in 5k bins, containing approximately 500 nuclei. PDX-P bulk: tumor tissue from CRC 2; PDX-P0,1,2,3 bulk: the zero, first, second and third-generation PDX (see the Methods). A consistent duplication is demonstrated in Chr. 6,7,8 for CRC 2 PDX-P0, P1, P2 and P3 bulk, which is shown as a fuzzy duplication in CRC 2 PDX-P bulk. In addition, an additional duplication is detected in Chr 2.

The successful rate of msCNVS library construction for these CTCs enriched from the two NSCLC patients and one CRC was acceptable. Here for NSCLC 1, 13 of 15 CTCs analyzed were qualified; for NSCLC 2, 19 of 20 CTCs qualified; and for CRC 1, 11 of 24 CTCs qualified (Fig 5c,d e and Extended Data Fig. 10a, b, c, d, e). However, the uniformity of sequences captured in library (reflected by NrcD values) for some batches of CTCs (ex. NSCLC 2, Fig. 5d and Extended Data Fig. 10c) were obviously better than other CTCs (NSCLC 1, CRC 1, Fig. 5 c,e and Extended Data Fig. 10a, b, d, e). Furthermore, for some batches of CTCs, the successful rate was low or all library constructions failed. Interestingly, msCNVS analysis for the WBCs from the batch of cells as of CTCs succeeded in less chance. From our experiences, whether a msCNVS library is qualified or not depends on the genome integrity of CTCs, which may be affected by the processes of CTC enrichment, isolation, identification, storage and transportation, in addition to the clinical therapy of the patient. It is likely that the genomes of these cells were degraded, or the cells were in process of apoptosis or other types of cell death.

We also employed msCNVS on single nuclei from a sum of 152 cells derived from PDX samples from patient CRC 2 (Extended Data Fig. 11). We compared single cell data to the corresponding bulk cell data (Fig. 5f). These nuclei all demonstrated CNAs, with similar patterns in Chr 6, 7, 8 between nuclei, which is consistent to an earlier reported result ^40^, However, a new mutation, a Chr 2 duplication, was detected in all single nuclei and the corresponding bulk nuclei in P3, which is supposed to be an outcome of cancer evolution. Therefore, multi-generation PDX is an interesting model for study of tumor evolution and for clinical treatment. msCNVS provides a useful tool for precise and efficient investigations of the CNV evolution in PDX.

### 2.6. Objective CNV calling based on SNrc peaks correlated to integer multiple copy number

As shown in the data above, we generated the SNrc density distribution collectively over the whole genome (Fig. 1c, e, Fig. 4c, d, Fig. 5b and Extended Data Fig. 3a, Extended Data Fig. 4e, Extended Data Fig. 10f, g, h). We found that the peak region (the wide range of the peak body) flanked by two valleys did not often exactly fall within the range of an integral multiple plus and minus 0.5 (ex. 1.5-2.5), nor did the valley often lie in the middle between two integral multiples (ex. 1.5 or 2.5). Therefore, the simple rounding adjustment for integer multiple CNV calling is not always objective. We analyzed the distribution of the major peaks demonstrated in the SNrc density distribution with the relationship to the copy number, so as to decide the integral multiple thresholds for CNV calling, as described in the Methods.

With this renovated protocol, there was only one major peak was identified, fitting the monomodal distribution, for the diploid cells GM12878 and Huvec as well as the healthy amniotic fluid cell cultures (Fig. 1e and Extended data Fig 3a). This almost exactly matched the 2-copy site on the SNrc distribution curves (gray label). The prominent major peak was similar in aneuploid cells of the amniotic fluid cell cultures (Extended data Fig. 3a), but there were some small major peaks close to 1-copy site (and/or 3-copy site) on the curves (blue or red labels), corresponding to segmental 1-copy, or 3-copy (or monosomy and trisomy of a whole chromosome), which overall fit the multimodal distribution. These callings gave the same output as that by the rounding adjustment used conventionally.

However, in some other samples, such as CTCs (Fig 5b and Extended data Fig. 3a, Extended data Fig.10f, g, h), the prominent major peak, obviously belonged to 2-copy (gray label), was in some degree switched up from the 2-copy region toward 3-copy region. And a smaller but major peak region located with a part of the peak region out of the range from 0.5-copy to 1.5-copy, which was determined as 1-copy (blue label). A similar scenario was observed for 3-copy (red labels). Meanwhile, a fuzzy peak zone (showing as a peak, but actually a fuzzy deletion zone) appeared between the 1-copy peak region and the 2-copy peak region (purple label in Fig 5b), and a fuzzy peak zone (a fuzzy duplication) was seen between the 2-copy peak region and the 3-copy peak region (pink label for PDX-P 5k bins, PDX-P2 5k bins in Extended Data Fig 11a,d) (pink label for PDX-P bulk 5k bins in Fig. 5f). From the data over this report, we noticed that for the same batch of samples, or the same nature of samples, the calling parameters (including thresholds) for different cells /samples were consistent, but for samples in different batches or with different properties, the parameters often varied. Taken together, for CNV calling with the new algorithm, we examined 5 types of calls, i.e., (a) <=1-copy, (b) fuzzy deletion (between 1- and 2-copy), (c) 2-copy, (d) fuzzy duplication (between 2- and 3-copy), (e) >=3-copy. However, additional types are easily extendable, such as 0-copy, or 4-copy +, and the fuzzy zones in between.

As we checked the CNV patterns of the same batch of single K562 cells with different resolutions or bins, obviously NrcD was more objective than CV value in reflection of the fluctuation of the SNrc profile (Extended data Fig. 4a, b, c, d). Under a given resolution, the lower the NrcD values indicates the better quality of the original data (Extended data Fig 3b). Under variants of resolution, a smaller NrcD values refers to a better quality of normalized count profile (Extended data Fig 4), which leads to higher confidence in CNV calling.

## 3. DISCUSSION

Next generation sequencing for CNV identification has been widely used to reveal insights into basic biology and inform clinical care decisions. scCNVS technology becomes increasingly important with the progress of single cell omics and precision medicine^1,41^. However, a series of problems exist in current conventional practice, such as the efficiency in cost and time, the requirement of preWGA with its associated consequences. msCNVS presented here is the first approach that implements pooling of multiple single cells individually pre-barcoded for CNV library construction, using off-the-shelf reagents and instruments and independent of preWGA (Supplementary File 2). Importantly, each input single cell is traceable from the sequencing data derived from the barcoded library fragments, and no cross contamination among single cells is detectable during the procedure. Moreover, with as low as 1Gb raw data per cell, CNV/CNA at 0.66 Mb resolution or 5k bins are consistently detectable; with deeper sequencing reads, or more cells (micro-bulk) such as the case of PGT-A, much higher resolution is achievable.

When msCNVS data of multiple single cells from the same cluster are merged in-silicon, the achievable CNV resolution is significantly raised. The reliability of msCNVS has been validated using single cells from 6 cases of amniotic fluid cell cultures. In these cells, the segmental and chromosomal trisomies and the deletions detected by msCNVS were confirmed by G-band chromosome karyotyping and CMA. In an application, apparent CNV heterogeneity, or mosaicism was detected in a discarded blastocyst, similar to what has been discovered by early reports ^2,39,42^. The analyses of CTCs and single nuclei from PDX tissue further support the efficiency and quality of the technique, demonstrating its power as a practical tool for clinical CNV detection, particularly in such situations as reproductive genetics and precision cancer therapy.

CNV calling from sequencing data has been a long-time challenge. In bulk CNV-seq, as we detected in the bulk cell sample of a CRC tissue (PDX-P 500 in Fig. 5f), ambiguous copy number values often exist, which is usually regarded as cellular heterogeneity of CNV^43^. However, in scCNS-seq for a given single cell, ambiguous copy number values also often show up. Such cases are difficult to explain because theoretically no heterogeneity is possible in single cell, which most likely are caused by cell-free DNA contamination when a single cell is isolated. Currently the rounding adjustment for integer multiples ^44^ is applied in multiple studies ^45,46^ to determine chromosomal or segmental CNVs. Although this rounding adjustment strategy may generally perform well for the ideal scenarios in scCNV-seq, in many cases the peak regions of the SNrc density distribution corresponding to 1-copy, 3-copy, or even 2-copy are out of the presumed ranges, and needed to be adjusted for integer copy numbers. To help solve this problem, this study also proposes a new strategy to define the major peaks with their boundaries associated with different integer copy number, based upon the monomodal (complete diploid) or multimodal (aneuploid) distribution of genome-wide SNrc density. This algorithm greatly improves the accuracy and objectivity of CNV calling.

As a method of low-pass sequencing, like the two recently reported DLP methods^20,47^, msCNVS gives the result of quantification of sparse sequencing reads over the whole genome. In addition, msCNVS does not efficiently co-detect copy neutral alterations such as SV (e.g., balanced translocations and inversions), short insertions and deletions (indels) and subtle SNVs. However, with SNV calling on the available sequencing data obtained in msCNVS, it is feasible to distinguish CNVs from copy-neutral losses of heterozygosity (CN-LOH) or UPD (uniparental disomy) in a relatively large scale ^13–16^. Interestingly, we find that silco-merged sequencing data of multiple cells, even of a few individual cells, greatly improves the resolution for CNV detection for a given clone or cluster of cells, as observed in other scenarios^20,47^.

The limiting step in the protocol of the current msCNVS method is the delivery of each single cell into a well, in addition to the initial barcoding of each cell within a restricted volume. Manual operation with a fine pipette and multichannel pipette, or mouth-operated micro-capillary pipette, fulfills the basic requirement when the cell number is limited^48^. Micromanipulation should further improve the cell pickup process^47^. When the purpose is to analyze a large number of cells or many biopsies in parallel, multiplex indexed pools can be combined (barcodes ×i5 ×i7) to elevate the throughput of msCNVS. A computer-controlled automatic system for single cell isolation, dispensing, and reagent delivery will further improve the efficiency^49^. By nature, msCNVS is easy to integrate with a microfluidic system such as droplet emulsification or microwell delivery, enabling a high throughput process with limited steps at the beginning for analyzing a large number of cells.

Correspondingly, the critical step for clinical application of msCNVS is the identification and isolation of the target cells from biopsies, such as the isolation of CTCs and MRDs of malignant tissue, trophoblast cells from preimplantation embryos or other fetal cells from pregnant women. The viability of the cells or the genome integrity of the nuclei is important for the success of msCNVS. Fortunately, there has been significant progress in recent research and clinical practice for enrichment and isolation of the target cells for PGT, NIPT and analysis of liquid biopsy of cancer^49–51^.

Overall, msCNVS is a robust method allowing for accurate CNV detection in single cells. msCNVS detects CNV with lower cost and higher efficiency by orders of magnitude compared with the conventional preWGA-based scCNV-seq methods^1,13–16,18,19^. Clinically, it outmatches the current clinical methods, such as G-band karyotyping and CMA, for fetal aneuploid analysis and embryo PGT-A. Meanwhile, CNV detection in CTCs by msCNVS provides genome-wide, large-scale mutation information, potentially guiding cancer precision medicine for early screening, monitoring, and therapy selection. Thus, msCNVS represents a valuable tool for clinical characterization of cellular CNV profile and heterogeneity at single cell level.

## Funding

National Nature Science Foundation of China (32071452)

National Nature Science Foundation of China (81770173)

Guangdong Major Basic Cultivation Project (2018B030308004)

Guangdong Natural Science Foundation Major Projects of Basic and Applied Basic Research (2019B1515120033)

Open Fund Programs of·Shenzhen Bar Laboratory (SZBL2020090501003)

Science and Technology Projects in Guangzhou (SL2022A04J01388)

## Author contributions

Conceptualization: Xinghua Pan

Methodology: Xinghua Pan, Guanchuan Lin, Bin Peng, Zhanying Dong, Mengchang Xu Investigation: Guanchuan Lin, Bin Peng, Zhanying Dong, Caimeng Chen, Mengchang Xu, Jingyu Gao, Jie Yu, Changtai Xiao, Linlin Wang, Liyao Mai, Yulong Zhang, Yuanfang Lu

Sample acquisition: Jie Yu, Bei Jia, Chen Luo, Rui Hua, Yuanqiao He, Yali Song, Mei Zhong, Song Quan

Visualization: Guanchuan Lin, Bin Peng, Zhanying Dong, Caimeng Chen, Mengchang Xu, Jingyu Gao, Changtai Xiao

Funding acquisition: Xinghua Pan Project administration: Xinghua Pan Supervision: Xinghua Pan, Hao Zhu,

Writing – original draft: Guanchuan Lin, Xinghua Pan

Writing – review & editing: Xinghua Pan, Hao Zhu, Sadie Marjani, Sherman Weissman, Weimin Zhang

## Data and materials availability

The public data used in this study is available in ENCODE project (ENCSR025GPQ, K562 bulk data); American National Library of Medicine(SRR10965088, GM12878 bulk data); NCBI Sequence Read Archive database (accession no. SRP052908, Huvec and of eMDA, MALBAC and MDA as well as HT-29 of DOP-PCR, MALBAC and MDA)

Data generated from this study have been deposited to the PRJCA018211. https://ngdc.cncb.ac.cn/

## Supplementary Materials

### Materials and Methods

#### Ethics

The human and mouse cell lines were stored at Southern Medical University. Human tissues including fetuses and embryos (blastocysts) and the related study were approved by Nanfang Hospital, Southern Medical University, with ethical number: NFEC-2019-194. The human samples were collected by registered professionals with the informed consent of the sample’s owners. The CTCs and cancer study were additionally approved by the Ethics Committee at the 2nd Affiliated Hospital, School of Medicine, Zhejiang University (2021-064) and the Ethics Committee at Sir Run Run Shaw Hospital, College of Medicine, Zhejiang University (20210323-41).

#### Propagation and single cell delivery for cell lines

Human cell lines, including K562, Jurkat, and HeLa, were cultured and passaged with RPMI 1640 Medium (Gibco, cat. no. C11875500CP). HeLa S3 cells were cultured and passaged with Ham’s F-12K Medium (CELLCOOK, cat. no. CM2004). HT-29 cells were cultured and passaged with McCoy’s 5A medium containing Alanyl-glutamine and HEPES (Procell, cat. no. PM150717). All media for the three cell lines above were supplemented with 10% FBS (HyClone, cat. no. SV30208.02). GM12878 cells were cultured and passaged with RPMI 1640 Medium, supplemented with 15% FBS. Huvec cells were cultured and passaged with Endothelial Cell Culture Medium (iCell Bioscience Inc, cat. PriMed-iCell-002) containing 93% ECM, 5% FBS, 1% Endothelial Cell Growth Supplement, and 1% Penicillin-Streptomycin. In addition, mouse cell line C2C12 cells were cultured and passaged with DMEM (Gibco, cat. no. C11995500BT), supplemented with 10% FBS. All cell lines above were cultured at 37°C with 5% CO_2_^52^. When Huvec cells became confluent^18^, they were washed twice with PBS, detached by 0.25% Trypsin-EDTA (1×) (Gibco, cat. no. 25200056), and centrifuged at 800 rpm for 3 min. Then, the supernatant was discarded, and the cell pellet was resuspended. All cell lines above tested negative for mycoplasma. Ten microliters of resuspended cells at a concentration of 1×10^6^ cells/ml were added into 1 ml precooled PBS with 10% FBS in a 6-well culture plate (NEST, 703001). Under an inverted microscope (Nikon, ECLIPSE Ts2), single cells were captured using 0.1-10 μl tips (Thermo Fisher Scientific, TFLR102-10-Q) and transferred into 8 stripe tubes (Axygen, PCR-02CP-C), one cell into each tube.

#### Amniotic fluid cell culture, G-band karyotyping analysis and chromosome microarray analysis (CMA)

Amniotic fluid samples were set up for cell culture following the standard protocols ^53^. Cells were cultivated in AmnioGrow Plus medium (Biowest, cat. no. 1083321) in a 37°C with 5% CO_2_ for 8-9 days and chromosome preparations were G-banded using trypsin-Giemsa staining. A minimum of 10 metaphases at the 400 to 500-band resolution level were routinely analyzed per specimen. Chromosome karyotype map scanning and acquisition were done using an automatic metaphase chromosome analysis system (MetaSystems, Göttingen, Germany). Two senior qualified laboratory technicians independently assessed karyotypes^54^.

For CMA, DNA was extracted from amniotic fluid samples according to standard protocols and microarrays were performed according to the manufacturer’s protocol with training provided by the industry donors of the microarray kits and reagents (Agilent Technologies and Affymetrix)^54,55^.

#### Single cell isolation and collection from discarded blastocysts

Modified from a reference protocol^56^, a 20 μl droplet of protease (QIAGEN, cat. no. 19155, dry powder 7.5 U, H_2_O or PBS, dilute to 1 U/ ml) was added into a well of a 35 mm dish, 60mm dish or a six-well plate. A discarded blastocyst (rated as 4BC) was transferred into the 20 μl droplet and incubated at 37°C for 15-30 min to digest the zona pellucida. The group of blastomere cells, partially aggregated and partially scattered, with digested zona pellucida were transferred to 20 μl Trypsin-EDTA solution, containing 0.25% trypsin and 0.02% EDTA (without phenol red, Ca^2+^ and Mg^2+^) and DNase I (working Con.=20 mg/l), reacted at 37°C for 5min. The cells were then transferred back to the 20 μl protease and incubated at 37°C for 10min. A 10 μl low-adsorption pipette tip with a filter was utilized to aspirate and disperse the cells. Finally, each cell was aspirated and dropped into a PCR tube for library construction.

#### Circulating tumor cells (CTC) isolation and identification

CTCs from peripheral blood were enriched by microfluidic HB CTC-Chip (Herringbone CTC chip), and identified by CTC EMT immunofluorescence staining kit for epithelial-mesenchymal subtyping, slightly modified from^51^. Specifically, the microfluidic chip chamber was functionalized with a combination of biotin-conjugated EpCAM (ThermoFisher, eBioscience) and CSV (Abnova), a coupled capture agent. The CTCs captured on the chip were stained with a cocktail of anti-Pan Cytokeratin monoclonal antibody (PanCK, AE1/AE3) (Alexa Fluor 488, ThermoFisher, eBioscience) and anti-vimentin monoclonal antibody (VIM, Alexa Fluor 488, CST). This was to collectively and efficiently identify epithelial (PanCK+), mesenchymal(VIM+) and epithelial-to-mesenchymal (PanCK+/VIM+) subtypes of CTCs based on epithelial mesenchymal transformation (EMT) mechanism. This was assisted by dual staining of PE anti-CD45 (Exbio, cat. No. 1P-222-T100) and PE anti-CD14 (CD45+/CD14+, Biolegend, cat. no. 301805) to determine white blood cells (WBC). DAPI (Hoechst 33342, ThermoFisher, cat. no. 62249) was used to stain all cell nuclei of CTC and WBC.

An automated BioView Duet scanning system, Allegro Plus (BioView, Rehovot, Israel), was utilized for HB CTC-Chip scanning and CTC identification. Positive staining for cytokeratin PanCK or VIM or both, without CD45 and CD14 staining, were identified as potential CTCs. Presumptive CTCs manually confirmed under 20× magnification using an automated inverted fluorescence microscope (Eclipse Ti, Nikon). WBCs were those cells with positive CD45 or CD14 staining. Single cells were isolated under 20× microscopic visualization using Nikon TI-FLC-E with NARISHIGE (MN-4) single cell micromanipulator.

#### PDX (Patient-Derived tumor Xenograft) sample preparation

We generated PDX lines following the protocol^57^ and supported by Nanchang Royo Biotech Co. Ltd. Five generations of PDX were collected: the original tumor tissue from the patient (P); the initial generation of PDX (P0) growing in a mouse inoculated with a piece of tumor tissue from the patient (P); the following generations of PDX (P1, P2, P3) each growing in a mouse transplanted from a piece of a graft tissue from the immediate early generation. The nuclei preparation was referred to Demonstrated Protocol CG000393 by 10x Genomics. A sample of PDX tissue approximately 0.2 g was cut into pieces in liquid nitrogen and transferred into a 15 ml centrifuge tube (Corning, cat. no. 430790). After incubating at 37°C until the tissue melted, 2 ml lysis buffer, including 100 µl of 1 M Tris-HCl (pH 7.4) (Sigma-Aldrich, cat. no. T2194), 20 µl of 5 M NaCl (Sigma-Aldrich, cat. no. 59222C), 30 µl of 1M MgCl_2_ (Sigma-Aldrich, cat. no. M1028), 100 µl of 10% IGEPAL ^®^ CA-630 (Sigma-Aldrich, cat. no. I3021), 9.75 ml of nuclease-free water (ThermoFisher, cat. no. AM9930) was added to the tissue for 6 min on ice. A serological pipette (NEST, cat. no. 318314) was used to pipette and mix during the lysis. Then 2ml of 1% BSA+PBS was added and the sample was centrifuged at 500 g for 5 min at 4°C. After removing the supernatant without disrupting the pellet, 1 ml of 1% BSA+PBS was added, and the pellet was gently resuspended. At the end, lysis efficiency and viability were assessed by staining the cells with trypan blue and monitored with the Countess II Automated Cell Counter (Thermo Fisher Scientific, cat. no. AMQAX1000). If a high fraction of viable cells was still present, the sample was centrifuged at 500 g for 5 min at 4°C and the lysis process was repeated. Finally, a 40 µm cell strainer was utilized to remove cellular debris if there was too much, and single nuclei capture was carried out similarly to the method described above in the section “Propagation and single cell isolation from cell lines”

#### msCNVS library generation

Before msCNVS library generation, the oligonucleotides associated with Tn5 transposase, which was utilized in this method, were specifically designed and optimized for efficient tagmentation and compatibility with the Illumina platform. The Tn5 adapter sequences of libraries consisted of three parts: oligo A, oligo B and oligo C. The oligo C was synthesized with a 5’ phosphorylated ME (Mosaic End) sequence and is partially complementary with both oligo A and B (Supplementary file 1). These oligos were coupled by annealing 4μl of each of two oligos (A or B with C, 100 μM) in a tube with 4μl of T4 ligation buffer and 8 µl of ddH_2_O, heating and cooling down to form double-stranded P5Tn5 adapter (formed by oligos A and C, with barcodes) and P7Tn5 adaptor (formed by oligos B and C, without barcodes). Finally, Tn5 transposase was added to assemble with Tn5 adapters (P5Tn5 adapter and P7Tn5 adapter) and form the customized active transposome. To generate the transposome, the Tn5 adapters were first diluted to a final concentration of 10 μM. Then, 1μl each of the two Tn5 adapters, 1 μl of 10×TPS buffer (Robustnique, cat. no. B0222), 2 μl of Tn5 transposase (1 U/ml, BGI, cat. no. 06R13) with 5 μl of dd H_2_O were mixed for incubating in a dry bath at 37°C for 30 min. The assembled Tn5 transposome was used for library construction or stored at −20°C.

As shown in Fig. 1a, Tn5 transposome was built of Tn5 transposase and the two Tn5 adapters. containing the newly designed 8-bp cell barcode in the Tn5P5 adapter. The two Tn5 adapters give PCR handles for the capturing 5’-end and 7’-end in PCR, which allows the common index and the cell-specific adapter sequences to be conjugated in the library by PCR, after sample pooling in one tube. The barcodes × P5 indexes × P7 indexes results in a B×i5×i7 combinations that enable simultaneous multiple single cell sequencing of CNV pattern at medium-throughput scale compatible with the Illumina platform.

After the Tn5 transposome was prepared in advance, cell lysis was performed before tagmentation. One microliter of diluted protease (Qiagen, cat. no. 19155, 7.5AU diluted with 7 ml H_2_O) was added to form a total 3 μl reaction including 1μl containing a single cell and 1 μl H_2_O. The mixture was incubated at 50°C for 1 h, with intermediate flicking of the tube at 15 min intervals followed by a brief spin down. After that, the protease was inactivated at 70°C for 30 min.

For tagmentation, lysis products were briefly centrifuged and then 1μl 5×LM buffer (Robustnique, cat. no. B0221) as well as 1μl of Tn5 transposome were added and incubated at 55°C for 20min. After that, 0.5µl of Exo 1 (Thermo Fisher, cat. no. EN0581) was added to make a 5 μl mixture of each tube, which was then incubated at 52°C for 20min, and then inactivated at 85°C for 15min. Finally, tagmentation was terminated by adding 1µl NT buffer (BGI, cat. no. 1000007876) or 0.2% SDS at 65°C for 30 min.

All tubes of the tagmentation products were pooled into a new EP tube and purified by DNA Concentration & Purification Kit (Zymo, cat. no. D4013). Then, 10 μl purified DNA, 2 μl primer index P7 (Vazyme, cat. no. TD202), 2 μl primer index P5 (Vazyme, cat. no. TD202), and 86 μl PCR master mix (Tsingke, cat. no. TSE101) were used for each reaction of PCR, with the thermocycler program set as follows: 72°C for 3 min, 98°C for 30s, 98°C for 15s, 60°C for 30 s, 72°C for 3 min (recycle step 3-5 for 22-30 cycles), 72°C for 5 min, finally hold at 4°C. The number of cycles was determined by the number of single cells in the mixed library, generally 27-28 cycles for 8 pooled single cells and 22-23 cycles for 48 pooled cells.

PCR products were purified by Zymo Concentration & Purification kit before 2% agarose E-gel electrophoresis separation and size selection. Alternatively, Ampure XP beads (Beckman Coulter, cat no. A63882) were used for size selection according to the manufacturer’s protocol. The fragment selection was set at 300-500 bp, alternatively 300 to 1000 bp was used. Finally, 20μl of ddH_2_O was added to elute the DNA.

Paired-end sequencing (PE150) with i5 and i7 indexes corresponding to P5 and P7 primers was performed. The library was sequenced in a whole lane or flowcell or combined with other libraries with the same structure. The desired amount of sequencing data was determined in advance.

#### msCNVS data analysis

To obtain the sequencing data of single cells, an in-house Python program was used to extract the sequencing library of each cell according to the barcode at the start of sequencing reads. Trimmomatic(v0.39)^58^ was used to remove the adapter sequences of each read and low quality reads. The sequence data was mapped to hg19 (human) or mm10 (mouse) using the Bowtie2^59^. Once mapped, the SAM files were transformed to BAM files using samtools^60^, and any reads not mapping to the reference genome were removed with parameters -bF 4. SortSam tools of Picard (2019, GitHub Repository https://broadinstitute.github.io/picard/, Broad Institute) was used to sort BAM files, and MarkDuplicates tools were used to remove duplicate reads with parameter REMOVE_DUPLICATES=true. Finally, BAM files were transformed into SAM files for downstream analysis.

Initially, we performed CNVs calling following the guidelines reported earlier^33^. The human genome was divided into bins of variable sizes based on the unique mapping of reads across the hg19 genome. GC content normalization was performed. After that, the median number of reads in each single bin was calculated from the SAM file. Cells with a median number of uniquely mapped reads less than 35 (in a bin) were removed. The counts within each bin were normalized according to GC content using Lowess smoothing as Normalized read count (Nrc). The Nrc data in each bin across all bins was further segmented using circular binary segmentation (CBS)^61^, becoming Segmented Nrc (SNrc).

Based on SNrc, rounding adjustment is a widely used method to yield integer copy number values and integral multiples is applied in different reports ^45,46^ to determine chromosomal or segmental monosomy, disomy or trisomy (51). Basically, a value with integer and fractional parts is rounded to the integer if the fractional part <0.5 or to the next integer if the fractional part >=0.5^44^. We found that the rounding adjustment generally performs well for the relatively ideal scenarios in scCNV-seq, but it is hard to make an objective calling in many other practical clinical cases.

To conduct CNV calling more objectively and accurately, we designed a strategy that exploits the distribution of SNrc density and identifies major peaks indicating of integer copy numbers, ie. segmental, chromosomal, or genome-wide monosomy (1-copy), disomy (2-copy), trisomy (3-copy), or higher ones when desired, along the genome (Extended data Fig 3. a). The strategy includes the following steps:

1. Computing the **SNrc Frequency Distribution** for all bins to unveil the distribution of the original SNrc values over the genome.
2. Transforming the SNrc Frequency Distribution to the **SNrc Density Distribution** to extract the integrated information of SNrc frequencies. For each bin (denoted by *bin_p*), the density of SNrc (denoted by *Den*(*SNrc_bin_p_*)) is computed as follows:

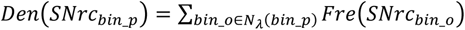

Here *bin_o* denotes any bin, *Fre*(*SNrc_bin_o_*) is the frequency of SNrc in *bin_o*, and *N_λ_*(*bin_p*) = {*bin_o*|*diff*(*bin_p*, *bin_o*) < *λ*} denotes the set of bins each has a copy number difference from the *bin_p* smaller than the threshold λ (this parameter is the upper limit of the copy number differences between two bins). Then, all 2-tuples (*SNrc_bin_p_*, *Den*(*SNrc_bin_p_*)) are projected to the coordinate system (X, Y).

1. Along the upper edge of the **SNrc Density Distribution (**the SNrc Density Distribution curve), a method of finding local maximum and minimum is used to scan SNrc values in

*Den*(*SNrc_bin_p_*) to identify major peaks on the SNrc Density Distribution:

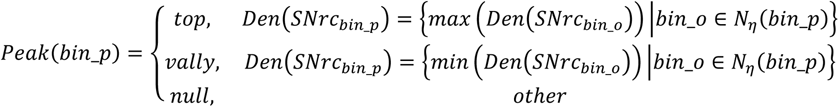

Here the parameter *η* determines the step size.

1. On the **SNrc Density Distribution,** the most prominent peaks near 1, 2, and 3 on the X-axis indicate 1-copy (monosomy), 2-copy (disomy), and 3-copy (trisomy), respectively. Each peak has two valleys at its two sides, and the regions between the valleys of two major peaks (e.g., the region between the valley of the 1-copy peak and the valley of the 2-copy peak) are called “fuzzy zones”. Fuzzy zone sizes can be adjusted empirically by setting *η* values according to the experimental references and CNV calling references reported in early studies.

Recognizing the influence of aneuploidy on CV, to accurately and objectively measure the experiment-derived signal deviation that may distort data quality, we evaluate whether the calling result of a cell (or sample) under a given bin number (resolution) is qualified, including whether the fluctuation of the SNrc signal for each copy (i.e., 1-copy, 2-copy, 3-copy, or more) in the genome-wide copy number profile (i.e., CNV pattern) collectively is too much and will make the CNV calling unreliable. To this end, we quantitatively define a NrcD (Nrc Deviation) value as follows to measure CNV calling quality:

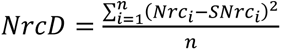

Here *i* refers to the *i-th* bin, and *n* represents the total number of bins. We empirically set the threshold of NrcD at 0.25 for msCNVS. When the NrcD value exceeds 0.25, the single cell CNV output is considered unqualified for reliable CNV calling under the given resolution or bin number. The bin number or resolution can be adjusted to meet the NrcD threshold, for example, by adjusting 5k bins downwards to 3k or 1.5k bins or upwards to 10k or 30k to increase the CNV resolution.

In addition to the NrcD, CV (Coefficient of Variation) is also calculated as the ratio of the standard deviation (SD) divided by the mean value of Nrc: CV=SD/mean. In the current study, in line with some early CNV reports ^18^, CV was used to evaluate the dispersion of Nrc values of all bins for a single cell (or, broadly speaking, sample) along the genome. Here, the Nrc value of each bin is used for CV analysis^18^. CV is affected by the amplitude and frequency of the segmental aneuploidies (or CNVs) of a cell (sample).

