## Supplemental Fig 1-11 and Table 1-2 for "msCNVS: medium throughput single cell copy number variation sequencing with barcoded library construction free of preamplification toward clinical implementation"

**Supplementary Text:**

**Supplementary file 1 Library and adaptors of msCNV:**

**msCNVS suquencing library**

5’**AATGATACGGCGACCACCGAGATCTACAC**(Index5)TCGTCGGCAGCGTCAGATGT GTATAAGAGACAGNNNNNNNNNNNN**AGATGTGTATAAGAGACAG**

-DNA insert-

CTGTCTCTTATACACATCTCCGAGCCCACGAGAC（**index7**）**ATCTCGTATGCCGTCTTCTGCTTG**-3’

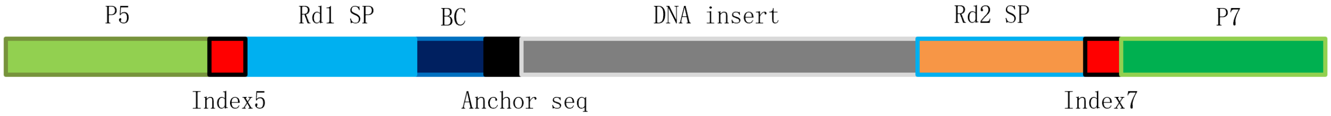


From 5’ to 3’: the P5 is used to anchor the library to the flowcell of Illumina NGS platforms. The index5 sequence is used to distinguish sample batches. Rd1 SP is the primer binding sequence for paired-end sequencing at one end. BC is the barcode sequence for single cell identification. The anchor sequencing is used to bind to Tn5 transposase for Tn5 transposome assembling. The gray part is the DNA insert (target). Rd2 SP is the other primer binding sequence of paired-end sequencing at the other end. The index7 is used to distinguish sample batch at the P7 end.

**Tn5 adaptor suquences**

Tn5 transposome includes a Tn5 transposase and two double strands of oligonucleotides, while one double strand oligos P5Tn5 adapter is annealed from oligo A and oligo C, and the other double strand oligos P7Tn5 adapter is annealed from oligo B and oligo C.

Oligo A:

| |
|---|
| TCGTCGGCAGCGTCAGATGTGTATAAGAGACAGNNNtcgccttaAGATGTGTATAAGAGACAG |
| TCGTCGGCAGCGTCAGATGTGTATAAGAGACAGNNNctagtacgAGATGTGTATAAGAGACAG |
| TCGTCGGCAGCGTCAGATGTGTATAAGAGACAGNNNttctgcctAGATGTGTATAAGAGACAG |
| TCGTCGGCAGCGTCAGATGTGTATAAGAGACAGNNNgctcaggaAGATGTGTATAAGAGACAG |
| TCGTCGGCAGCGTCAGATGTGTATAAGAGACAGNNNaggagtccAGATGTGTATAAGAGACAG |
| TCGTCGGCAGCGTCAGATGTGTATAAGAGACAGNNNcatgcctaAGATGTGTATAAGAGACAG |
| TCGTCGGCAGCGTCAGATGTGTATAAGAGACAGNNNgtagagagAGATGTGTATAAGAGACAG |
| TCGTCGGCAGCGTCAGATGTGTATAAGAGACAGNNNcagcctcgAGATGTGTATAAGAGACAG |
| TCGTCGGCAGCGTCAGATGTGTATAAGAGACAGNNNtgcctcttAGATGTGTATAAGAGACAG |
| TCGTCGGCAGCGTCAGATGTGTATAAGAGACAGNNNtcctctacAGATGTGTATAAGAGACAG |
| TCGTCGGCAGCGTCAGATGTGTATAAGAGACAGNNNtcatgagcAGATGTGTATAAGAGACAG |
| TCGTCGGCAGCGTCAGATGTGTATAAGAGACAGNNNcctgagatAGATGTGTATAAGAGACAG |
| TCGTCGGCAGCGTCAGATGTGTATAAGAGACAGNNNtagcgagtAGATGTGTATAAGAGACAG |
| TCGTCGGCAGCGTCAGATGTGTATAAGAGACAGNNNgtagctccAGATGTGTATAAGAGACAG |
| TCGTCGGCAGCGTCAGATGTGTATAAGAGACAGNNNtactacgcAGATGTGTATAAGAGACAG |
| TCGTCGGCAGCGTCAGATGTGTATAAGAGACAGNNNaggctccgAGATGTGTATAAGAGACAG |
| TCGTCGGCAGCGTCAGATGTGTATAAGAGACAGNNNgcagcgtaAGATGTGTATAAGAGACAG |
| TCGTCGGCAGCGTCAGATGTGTATAAGAGACAGNNNctgcgcatAGATGTGTATAAGAGACAG |
| TCGTCGGCAGCGTCAGATGTGTATAAGAGACAGNNNgagcgctaAGATGTGTATAAGAGACAG |
| TCGTCGGCAGCGTCAGATGTGTATAAGAGACAGNNNcgctcagtAGATGTGTATAAGAGACAG |
| TCGTCGGCAGCGTCAGATGTGTATAAGAGACAGNNNgtcttaggAGATGTGTATAAGAGACAG |
| TCGTCGGCAGCGTCAGATGTGTATAAGAGACAGNNNactgatcgAGATGTGTATAAGAGACAG |
| TCGTCGGCAGCGTCAGATGTGTATAAGAGACAGNNNtagctgcaAGATGTGTATAAGAGACAG |
| TCGTCGGCAGCGTCAGATGTGTATAAGAGACAGNNNgacgtcgaAGATGTGTATAAGAGACAG |
| TCGTCGGCAGCGTCAGATGTGTATAAGAGACAGNNNctctctatAGATGTGTATAAGAGACAG |
| TCGTCGGCAGCGTCAGATGTGTATAAGAGACAGNNNtatcctctAGATGTGTATAAGAGACAG |
| TCGTCGGCAGCGTCAGATGTGTATAAGAGACAGNNNgtaaggagAGATGTGTATAAGAGACAG |
| TCGTCGGCAGCGTCAGATGTGTATAAGAGACAGNNNactgcataAGATGTGTATAAGAGACAG |
| TCGTCGGCAGCGTCAGATGTGTATAAGAGACAGNNNaaggagtaAGATGTGTATAAGAGACAG |
| TCGTCGGCAGCGTCAGATGTGTATAAGAGACAGNNNctaagcctAGATGTGTATAAGAGACAG |
| TCGTCGGCAGCGTCAGATGTGTATAAGAGACAGNNNcgtctaatAGATGTGTATAAGAGACAG |
| TCGTCGGCAGCGTCAGATGTGTATAAGAGACAGNNNtctctccgAGATGTGTATAAGAGACAG |
| TCGTCGGCAGCGTCAGATGTGTATAAGAGACAGNNNtcgactagAGATGTGTATAAGAGACAG |
| TCGTCGGCAGCGTCAGATGTGTATAAGAGACAGNNNttctagctAGATGTGTATAAGAGACAG |
| TCGTCGGCAGCGTCAGATGTGTATAAGAGACAGNNNcctagagtAGATGTGTATAAGAGACAG |

| |
|---|
| TCGTCGGCAGCGTCAGATGTGTATAAGAGACAGNNNgcgtaagaAGATGTGTATAAGAGACAG |
| TCGTCGGCAGCGTCAGATGTGTATAAGAGACAGNNNctattaagAGATGTGTATAAGAGACAG |
| TCGTCGGCAGCGTCAGATGTGTATAAGAGACAGNNNaaggctatAGATGTGTATAAGAGACAG |
| TCGTCGGCAGCGTCAGATGTGTATAAGAGACAGNNNgagccttaAGATGTGTATAAGAGACAG |
| TCGTCGGCAGCGTCAGATGTGTATAAGAGACAGNNNttatgcgaAGATGTGTATAAGAGACAG |
| TCGTCGGCAGCGTCAGATGTGTATAAGAGACAGNNNtatagcctAGATGTGTATAAGAGACAG |
| TCGTCGGCAGCGTCAGATGTGTATAAGAGACAGNNNatagaggcAGATGTGTATAAGAGACAG |
| TCGTCGGCAGCGTCAGATGTGTATAAGAGACAGNNNcctatcctAGATGTGTATAAGAGACAG |
| TCGTCGGCAGCGTCAGATGTGTATAAGAGACAGNNNggctctgaAGATGTGTATAAGAGACAG |
| TCGTCGGCAGCGTCAGATGTGTATAAGAGACAGNNNaggcgaagAGATGTGTATAAGAGACAG |
| TCGTCGGCAGCGTCAGATGTGTATAAGAGACAGNNNtaatcttaAGATGTGTATAAGAGACAG |
| TCGTCGGCAGCGTCAGATGTGTATAAGAGACAGNNNcaggacgtAGATGTGTATAAGAGACAG |
| TCGTCGGCAGCGTCAGATGTGTATAAGAGACAGNNNgtactgacAGATGTGTATAAGAGACAG |

Oligo B:

GTCTCGTGGGCTCG**AGATGTGTATAAGAGACAG**

Oligo C:

5'PHOS-CTGTCTCTTATACACATCT

### Supplementary file 2:

| Time | Hands-on Time: Bench | | Hands-Off Time: Instrument | | Temperature | Stop&Storage |
|---|---|---|---|---|---|---|
| | Single cell delivery | variable | | | | |
| 0h | Cell lysis and DNA exposure | | | | | |
| | Reagent preparation | 10min | | | RT | |
| | Reagent loading | 10min | | | RT | |
| | | | Cell lysis/protein digestion | 20-40min | 50℃ | |
| | | | Enzyme deactivation | 30min | 70℃ | |
| 1.5h | Barcoding | | | | | |
| | Buffer loading | 10min | | | | |
| | Tn5 complex loading | 10min | | | | |
| 2h | | | Cell barcoding | 20min | 55℃ | |
| | Exo1 loading | 10min | | | | |
| | | | Excess oligos elimination | 20min | 52℃ | |
| | | | | 15min | 85℃ | |
| | SDS loading | 10min | | | | |
| 3h | | | Tn5 deactivation | 30min | 65℃ | |
| | Indexing | | | | | |
| | Samples pooling | 10min | | | | |
| | DNA purification | 10min | | | | |
| 3.5h | Index adaptor loading | 10min | | | | |
| | | | Library PCR | 90min | | |
| 5.5h | Library purification | 10min | | | | 4℃<24h, -20℃>24h |
| | Library pufication | | | | | |
| | DNA loading | 5min | | | | |
| | | | E-gel running | 30min | | |
| | Size selection on gel | 20min | | | | |
| 6.5h | Gel purification | 20min | | | | 4℃<24h, -20℃>24h |
| | | | DNA Quantification on Qubit | 5min | | |
| | Total | 145min | | 260min-280min | | |

### Figs. S1 to S11:

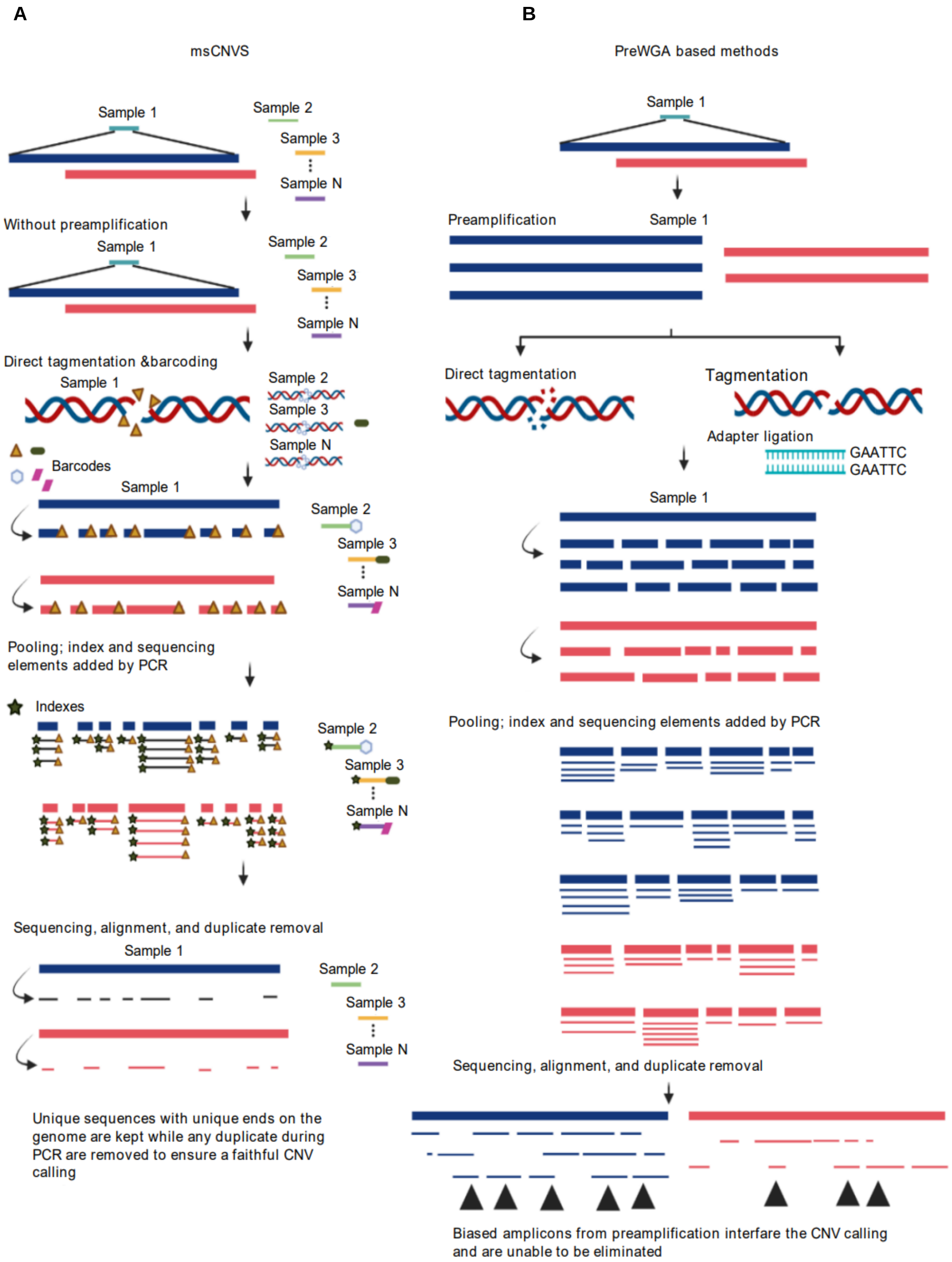

**Extended Data Fig. 1. Comparison of msCNVS with the scCNV-seq methods based on single cell pre-whole genome amplification (preWGA).**

**A. msCNVS. msCNVS labels the natural genome of each cell through an oligonucleotide-barcode contained in a special Tn5 transposome complex at the very beginning, followed by pooling of multiple single cells.** The CNV sequencing libraries of these cells/genomes in a single tube are then collectively constructed, and the batch index for the pooled samples is linked to the sequencing library by PCR. Without preWGA of the individual genomes, msCNVS ensures only the natural chromosomal sequences to be captured randomly each with a pair of unique ends of the insert. These unique ends, after sequencing, are based to retain the unique sequences and to remove any PCR-introduced duplicates. Subsequently a faithful calling of CNVs is achievable. Meanwhile, the cell-specific barcodes and batch indexes are used to trace the original identification of each single cell.

**B. Methods with preWGA.** Conventional methods require a independent procedure of preWGA for each single cell genome to be individually amplified. After preWGA, the amplicon of each single cell is then independently converted into a sequencing lbrary, which is also built with cell barcode and index. Because the library construction is on the preWGA amplicon, additional amplifictaion bias and sequence duplication introduced in the preWGA off the original sequences cannot be removed, which interferes the CNV calling.

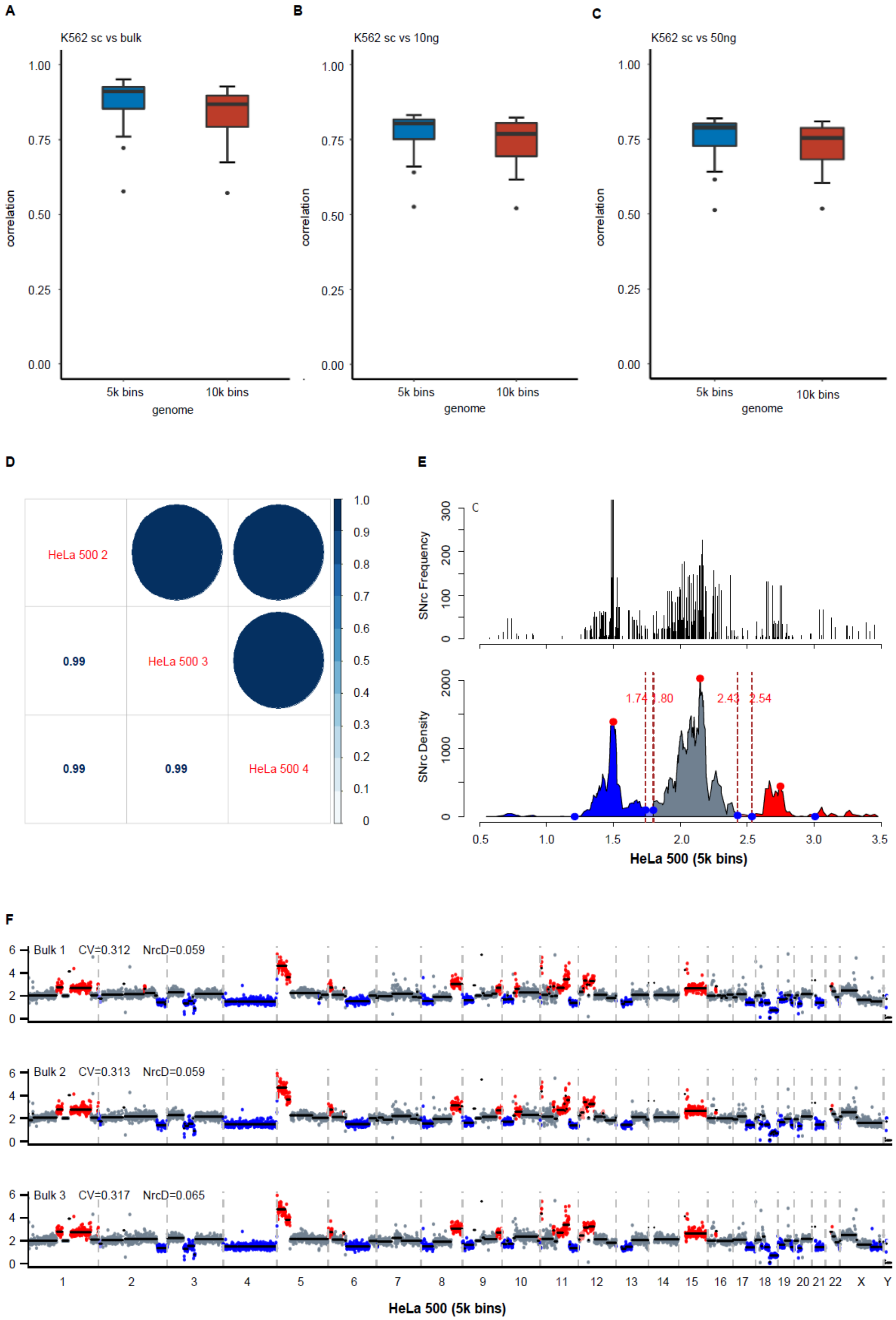
A
K562 sc vs bulk
correlation
1.00
0.75
0.50
0.25
0.00
5k bins
10k bins
genome
B
K562 sc vs 10ng
C
K562 sc vs 50ng
D
HeLa 500 2
HeLa 500 3
HeLa 500 4
0.99
0.99
0.99
1.0
0.9
0.8
0.7
0.6
0.5
0.4
0.3
0.2
0.1
0
E
SNrc Frequency
SNrc Density
1.74
1.80
2.43
2.54
HeLa 500 (5k bins)
F
Bulk 1 CV=0.312 NrcD=0.059
Bulk 2 CV=0.313 NrcD=0.059
Bulk 3 CV=0.317 NrcD=0.065
1 2 3 4 5 6 7 8 9 10 11 12 13 14 15 16 17 18 19 20 21 22 X Y
HeLa 500 (5k bins)

**Extended Data Fig. 2. Corelation of CNV profiles between single cells by msCNVS and bulk, and reproducibility of bulk cells by mCNVS.** Two resolutions apply: 0.66Mb (5k bins) and 0.33Mb (10k bins). The correlation as well as the CNVpattern plot are analyzed with ggplot (R).

**A. Between K562 single cells (n=70) by msCNVS and bulk data from ENCODE.**

**B. Between K562 single cells (n=70) by msCNVS and bulk data (10ng DNA) by msCNVS.**

**C. Between K562 single cells (n=70) by msCNVS and bulk data (50ng DNA) by msCNVS.**

**D. Correlations of Triplicates of Hela cells in bulk (n≈500 cells) by mCNVS (5k bins, 0.66Mb).**

**E. CNV calling based on SNrc Density Distribution and Frequency Distribution of merged triplicates of HeLa cells in bulk.**

**F. The CNV patterns of HeLa cells in bulk (n≈500 cells) by mCNVS (5k bins, 0.66Mb) with triplicates.**

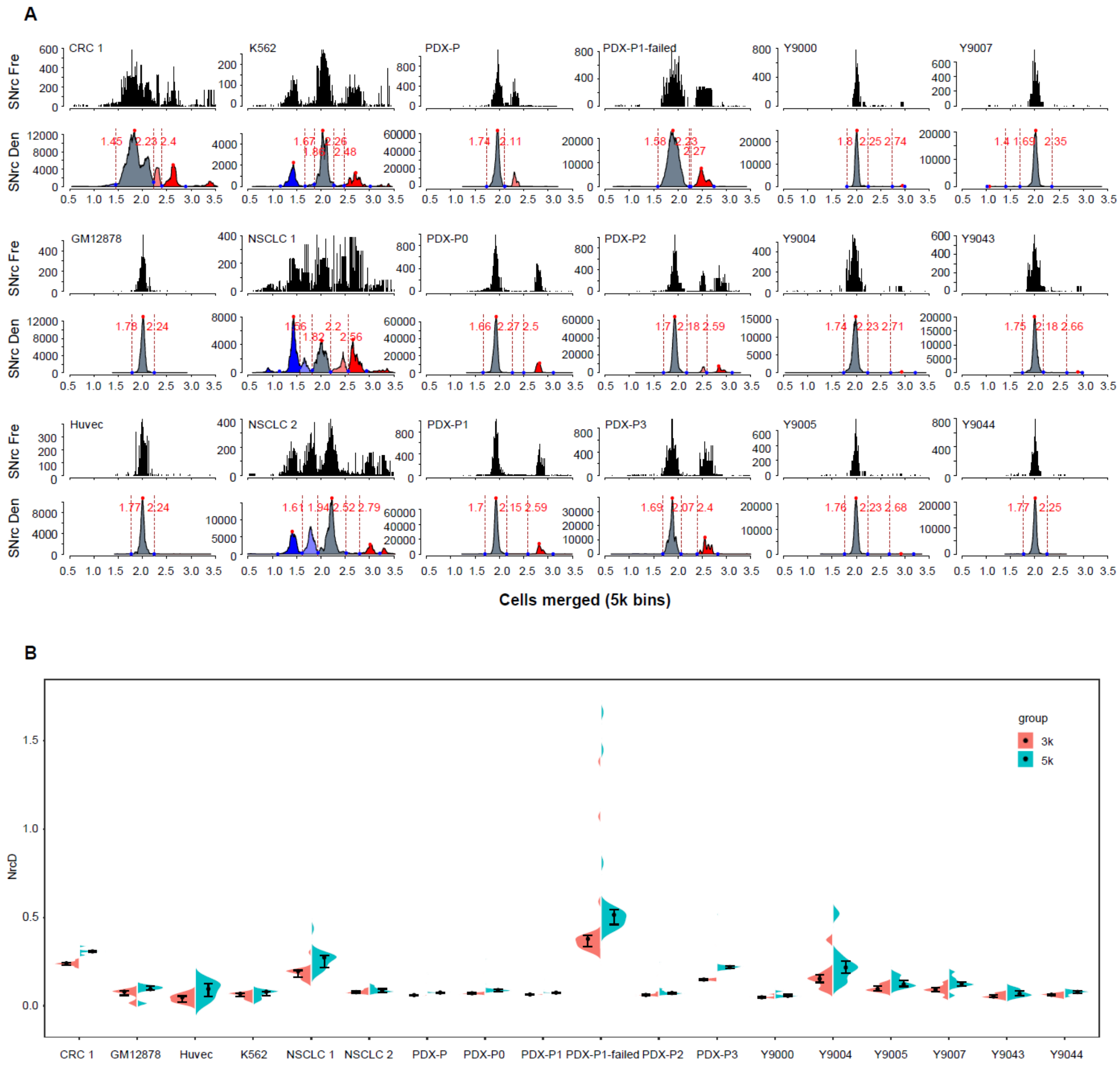


### Extended Data Fig 3

**A. SNrc Frequency Distribution and SNrc Density Distribution of merge data of single cells generated by msCNVS for different samples.** The merged data for single cells at the resolution of 5k bins are displayed for all samples.

"K562" refers to Fig 1 B, C & Extended Data Fig 4 C for K562.

"GM12878" refers to Fig 1 D, E for GM12878.

"Huvec" refers to Fig 2 A for Huvec.

"NSCLC 1" refers to Fig 5 C & Extended Data Fig 10 B for NSCLC 1;

"NSCLC 2" refers to Fig 5 D & Extended Data Fig 10 C for NSCLC 2;

"CRC 1" refers to Fig 5 E & Extended Data Fig 10 E for CRC1 CTCs;

"PDX-P" refers to Extended Data Fig 11 A for PDX-P.

"PDX-P0" refers to Extended Data Fig 11 B for PDX-P0.

"PDX-P1" refers to Extended Data Fig 11 C for PDX-P1.

"PDX-P1 failed" was displayed alone.

"PDX-P2" refers to Extended Data Fig 11 D for PDX-P2.

"PDX-P3" refers to Extended Data Fig 11 E for PDX-P3.

"Y9000" refers to Fig 3 D, G for Y9000.

"Y9005" refers to Fig 3 E, I for Y9005.

"Y9007" refers to Fig 3 F, K for Y9007.

"Y9004" refers to Extended Data Fig 8 A for Y9004.

"Y9043" refers to Extended Data Fig 8 B for Y9043.

"Y9044" refers to Extended Data Fig 8 C for Y9044.

**B. Comparison of the NrcD values of the CNV patterns at two resolutions (3k and 5k bins).**
In this study, it is empirically set the threshold of NrcD value at 0.25 to judge the qualification of the msCNVS data. When NrcD is higher than 0.25, the data of the single cell under the given resolution is regarded as unqualified for a faithful CNV calling.

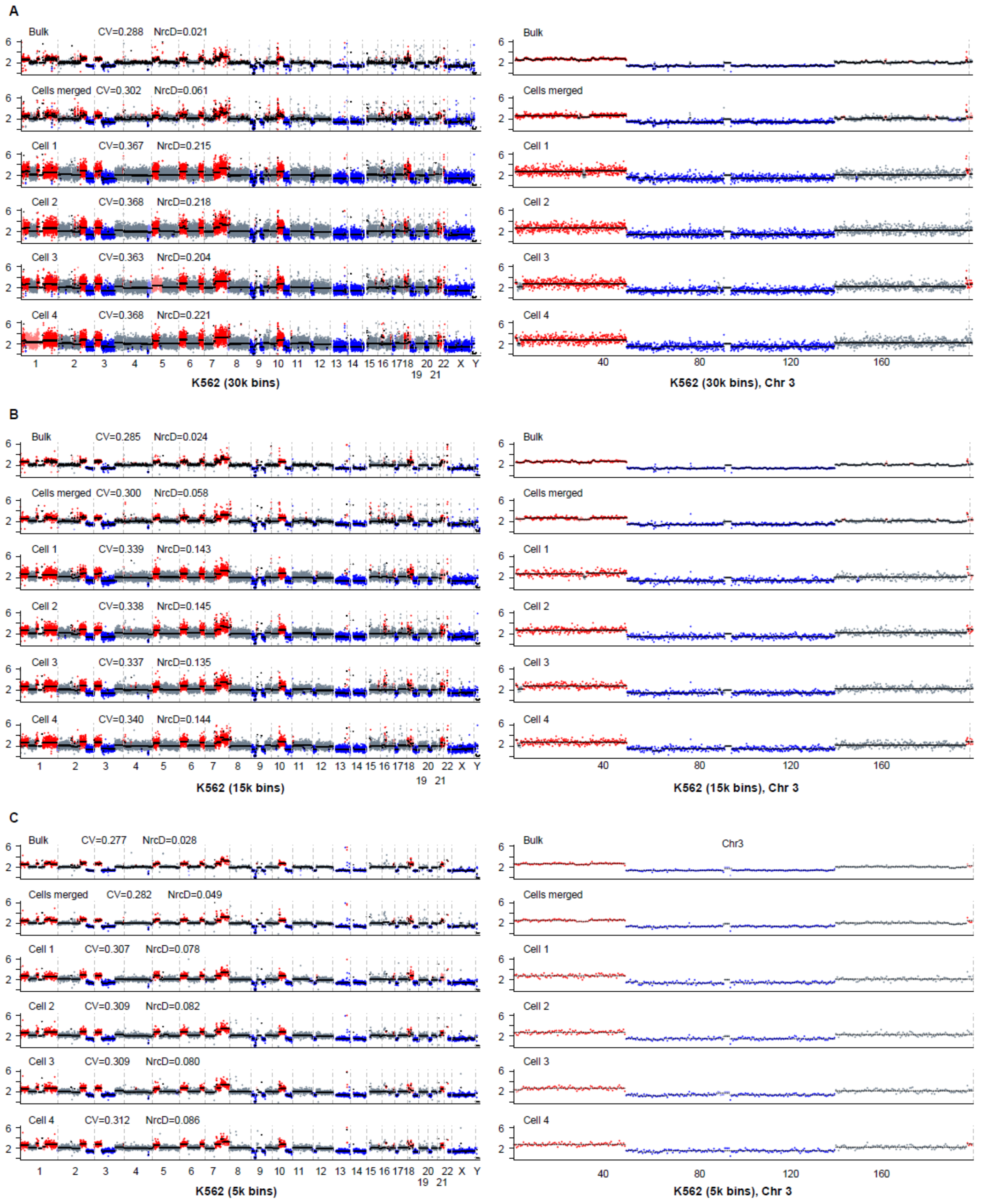
A
Bulk CV=0.288 NrcD=0.021
Cells merged CV=0.302 NrcD=0.061
Cell 1 CV=0.367 NrcD=0.215
Cell 2 CV=0.368 NrcD=0.218
Cell 3 CV=0.363 NrcD=0.204
Cell 4 CV=0.368 NrcD=0.221
K562 (30k bins)
K562 (30k bins), Chr 3
B
Bulk CV=0.285 NrcD=0.024
Cells merged CV=0.300 NrcD=0.058
Cell 1 CV=0.339 NrcD=0.143
Cell 2 CV=0.338 NrcD=0.145
Cell 3 CV=0.337 NrcD=0.135
Cell 4 CV=0.340 NrcD=0.144
K562 (15k bins)
K562 (15k bins), Chr 3
C
Bulk CV=0.277 NrcD=0.028
Cells merged CV=0.282 NrcD=0.049
Cell 1 CV=0.307 NrcD=0.078
Cell 2 CV=0.309 NrcD=0.082
Cell 3 CV=0.309 NrcD=0.080
Cell 4 CV=0.312 NrcD=0.086
Chr3
K562 (5k bins)
K562 (5k bins), Chr 3

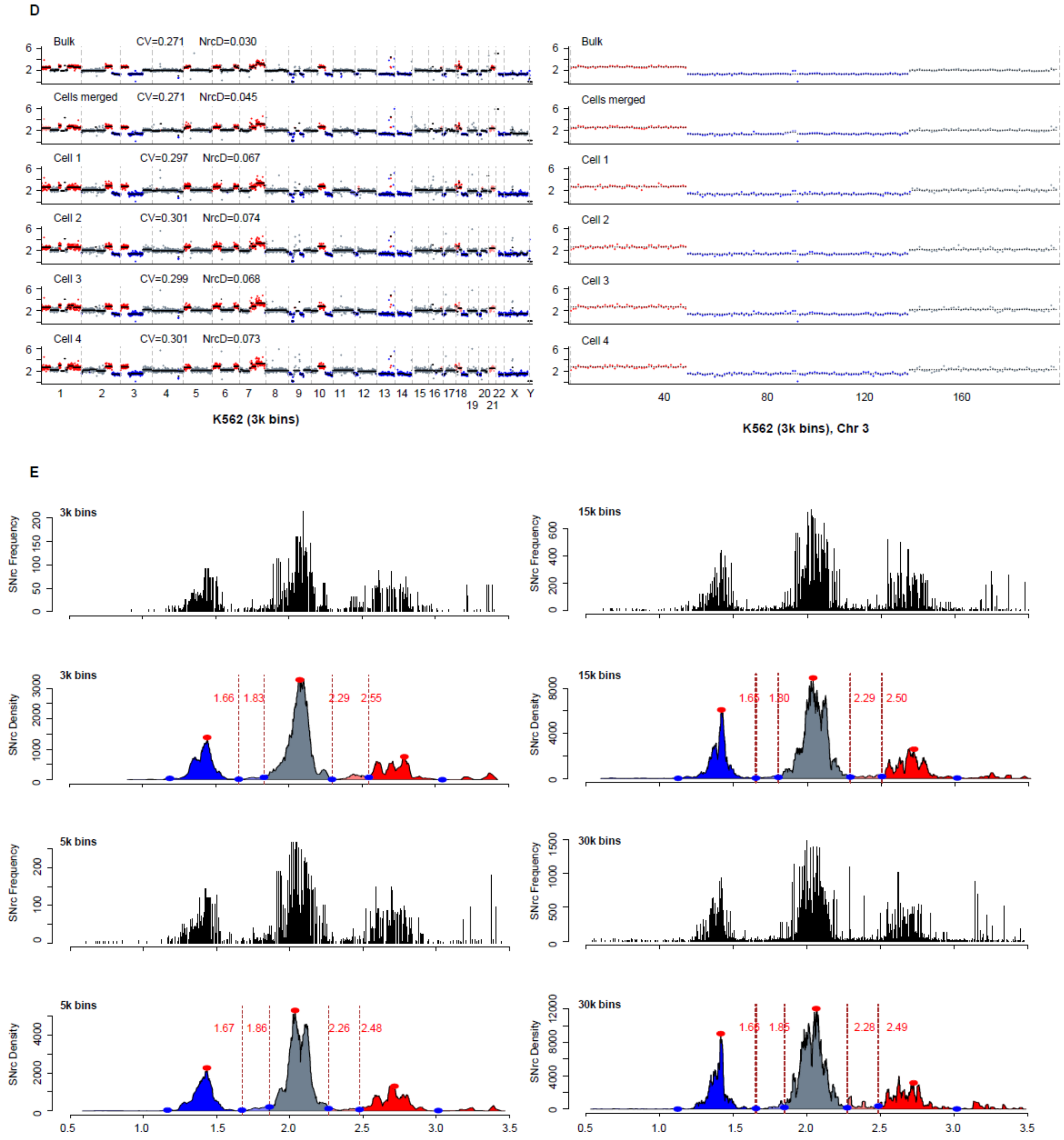


**Extended Data Fig. 4. CNV petterns of K562 single cells generated by msCNVS analyzed at different resolutions, showing different signal fluctuations.** The merged data of 70 single cells and 4 randomly chosen individual single cells are displayed in comparison to bulk data from ENCODE, visualized at 4 different resolutions. All samples (single cells and bulk) show a similar CNV patterns particualy at the large scale CNVs (ex. chromosome 3: segmental duplication, deletion and normal diploid, as showing on the right panel) among different resolutions. Overall, the signal fluctuation increase from bulk, merged sample, to single cells. However, at relatively higher resolution (A) versus lower resolution (D), more subtle (small) CNVs are detectable in higher resolution, with a higher signal fluctation, and vice versa. It is noted that the merged data of single cells shows little signal fluctuation similar to the bulk, and it also detects more small CNVs similar to the bulk, and that NrcD more faithfully reflects the signal fluctuation than CV (coefficient of variation) value.

Left: whole genome. Right: zoomed chromosome 3.

**A. 0.1 Mb (30,000 bins)**

**B. 0.2 Mb (15,000 bins)**

**C. 0.6 Mb (5,000 bins)**

**D. 1.0 Mb (3,000 bins)**

**E. SNrc Frequency Distribution and SNrc Density Distribution of merged data for K562 single cells at different resolutions.**

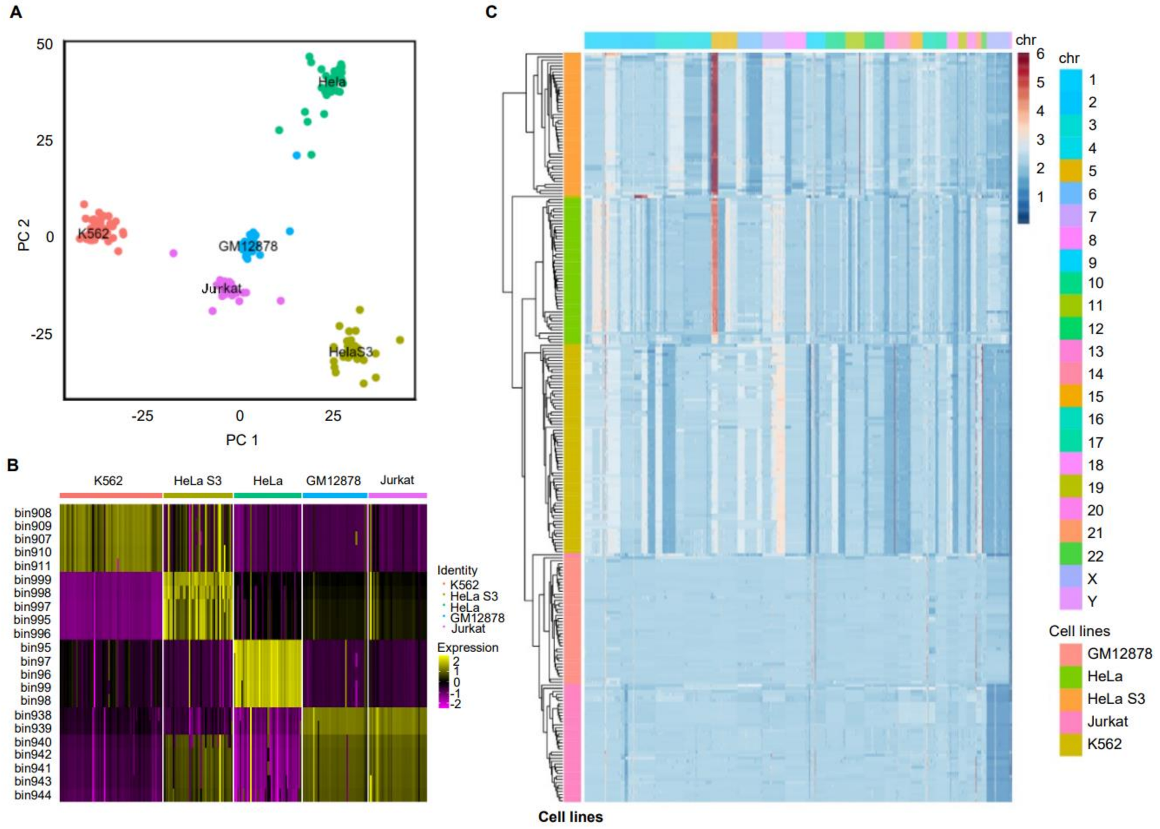


**Extended Data Fig. 5. Comparison of CNV heterogeneity among single cells for each of 5 human cell lines, measured by msCNVS, with their clusterings and features.** The cell numbers are as follows: K562 (n=70), GM12878 (n=48), Hela (n=48), Hela S3 (n=47), Jurkat(n=32).

**A. PCA clustering analysis. Single cells from different cell lines are gathered with the same cell line.**

**B. Different CNV regions among 5 human cell lines based on msCNVS analyzed on Seurat. The genome is divided into 5k bins (at 0.66Mb resolution).**

**C. Heatmap showing the similarity and heterogeneity within each of the cell lines, and the relationship among the cell lines based on the msCNVS pattern of each single cell.**

A

B

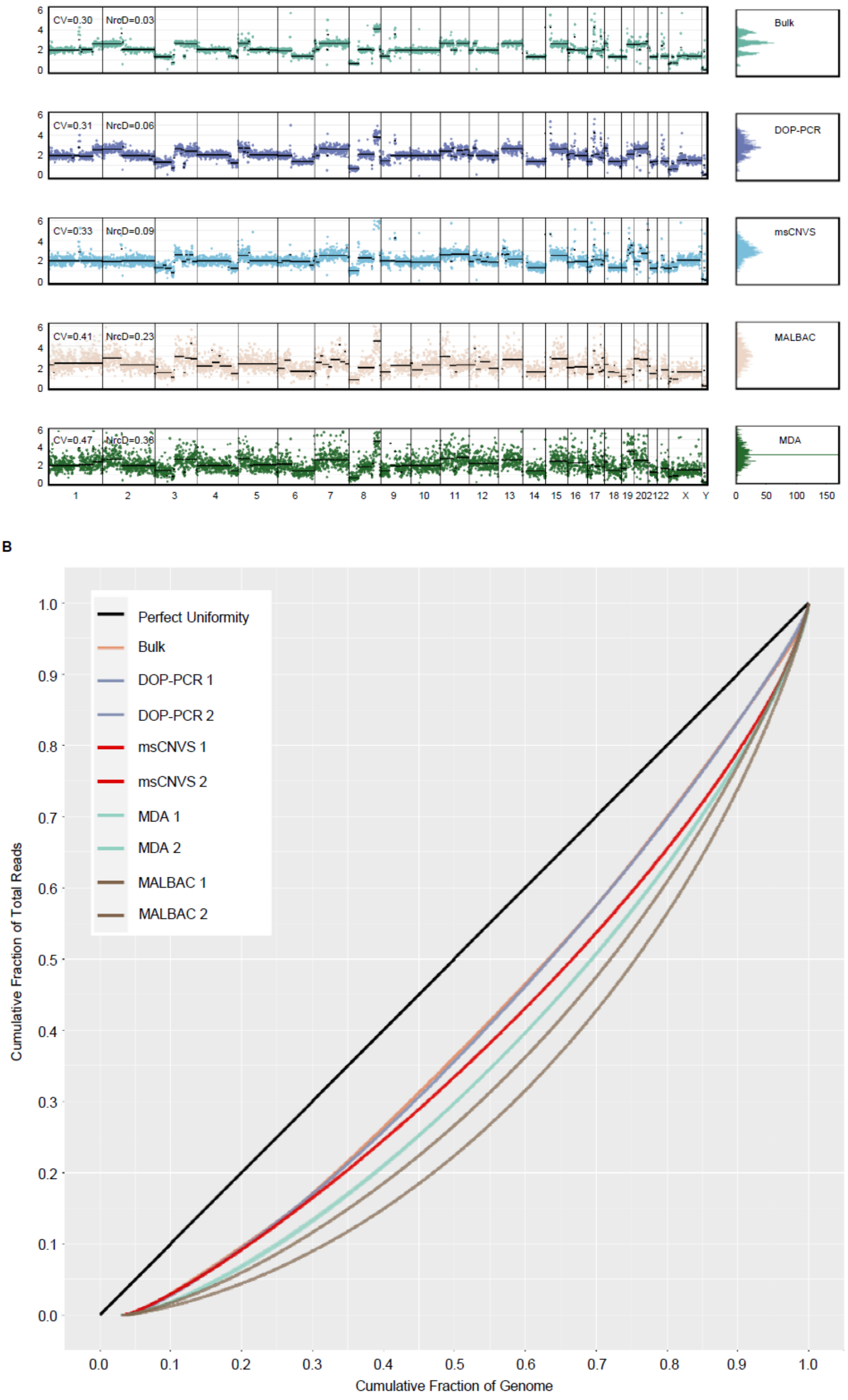

CV=0.30 NrcD=0.03
Bulk
CV=0.31 NrcD=0.06
DOP-PCR
CV=0.33 NrcD=0.09
msCNVS
CV=0.41 NrcD=0.23
MALBAC
CV=0.47 NrcD=0.36
MDA
1 2 3 4 5 6 7 8 9 10 11 12 13 14 15 16 17 18 19 20 21 22 X Y
0 50 100 150
Perfect Uniformity
Bulk
DOP-PCR 1
DOP-PCR 2
msCNVS 1
msCNVS 2
MDA 1
MDA 2
MALBAC 1
MALBAC 2
Cumulative Fraction of Total Reads
Cumulative Fraction of Genome

**Extended Data Fig. 6. Coverage uniformity of msCNVS in comparison to DOP-PCR, msCNVS, MALBAC and MAD for single cells of cell line HT-29.**

**A. Comparing the CNV patterns generated by 5 different methods. CV (coefficient of variation) and NrcD are applied to show the level of signal fluctation.** The data for bulk， DOP-PCR, MDA and MALBAC are obtained from public resouces (NCBI Sequence Read Archive database (accession no. SRP052908)). The msCNVS data is generated by msCNVS in this report, which is slightly different from the public database, being probably attributed to some new mutations in the HT-29 during passaging in culture.

**B. Lorenz curve of coverage uniformity of CNV patterns, of which two single cells are displayed for each of the 5 methods.**

A K562

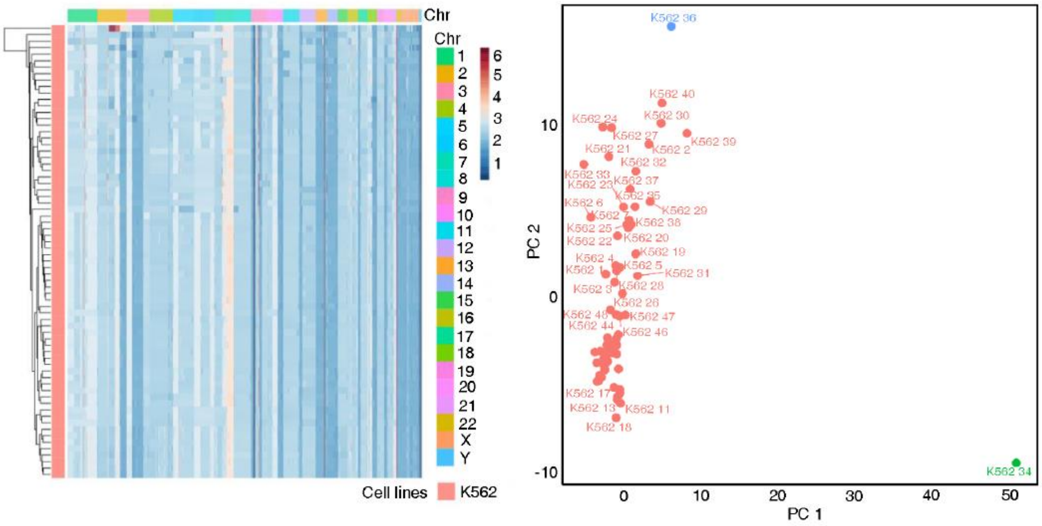


B HeLa

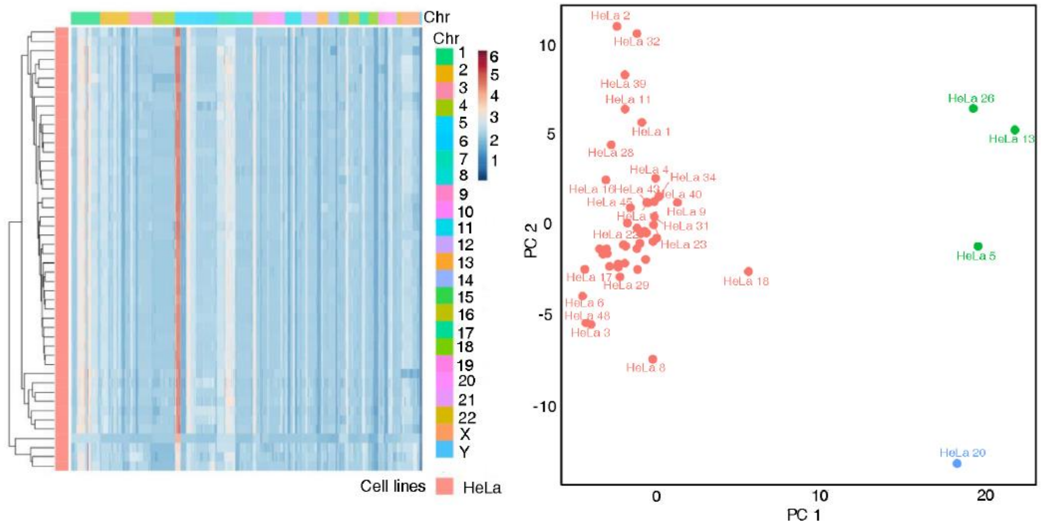


C HelaS3

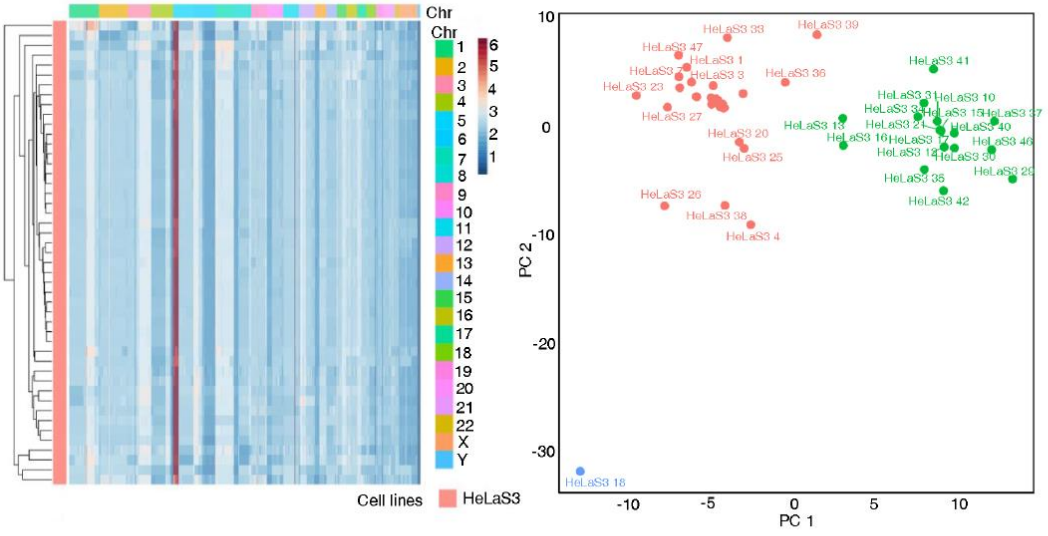

**Extended Data Fig. 7. CNV heterogeneity among single cells within each of the 3 cell lines, K526, Hela and Hela S3.** The CNV patterns are generated by msCNVS, and displayed by heatmap and PCA clustering.

**K526 cells (n=70).**

**Hela cells (n=48).**

**Hela S3 cells (n=47).**

A

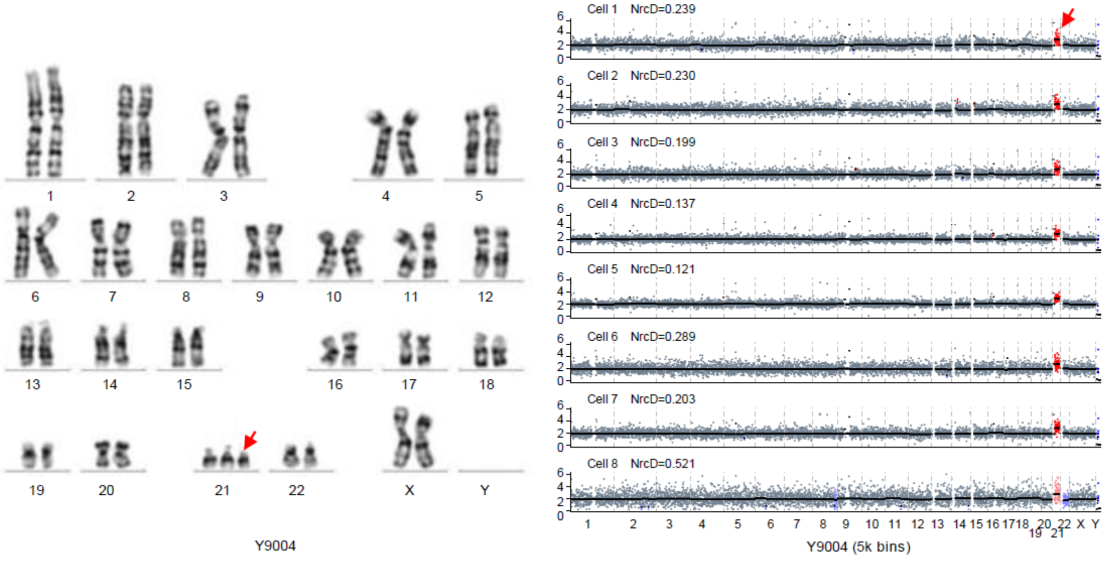
Y9004
Cell 1 NrcD=0.239
Cell 2 NrcD=0.230
Cell 3 NrcD=0.199
Cell 4 NrcD=0.137
Cell 5 NrcD=0.121
Cell 6 NrcD=0.289
Cell 7 NrcD=0.203
Cell 8 NrcD=0.521
Y9004 (5k bins)

B

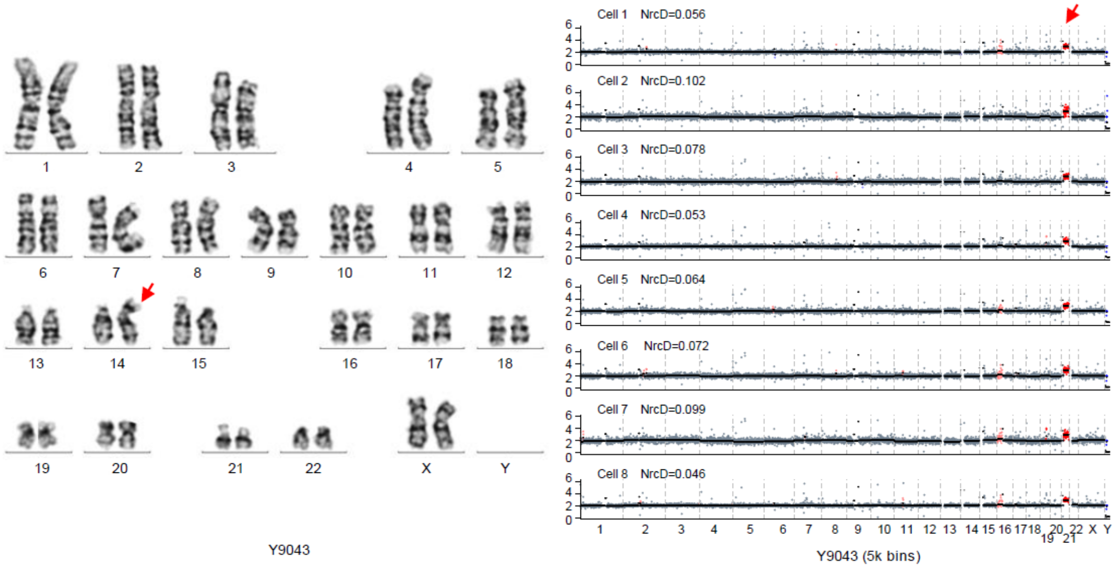
Y9043
Cell 1 NrcD=0.056
Cell 2 NrcD=0.102
Cell 3 NrcD=0.078
Cell 4 NrcD=0.053
Cell 5 NrcD=0.064
Cell 6 NrcD=0.072
Cell 7 NrcD=0.099
Cell 8 NrcD=0.046
Y9043 (5k bins)

C

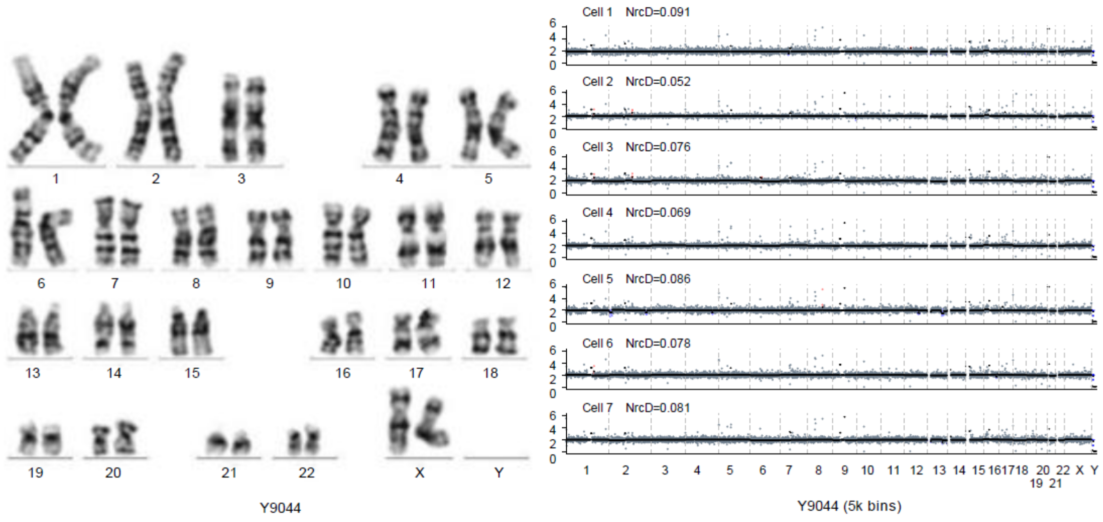
Y9044
Cell 1 NrcD=0.091
Cell 2 NrcD=0.052
Cell 3 NrcD=0.076
Cell 4 NrcD=0.069
Cell 5 NrcD=0.086
Cell 6 NrcD=0.078
Cell 7 NrcD=0.081
Y9044 (5k bins)

**Extended Data Fig. 8. Detection of trisomy 21 by msCNVS with single cells from the amniotic fluid cell culture from 2 patients.** msCNVS pattern (right) is consistent to G-banding kayotype (left).

**A. Standard trisomy 21 (Y9004, n=7/8).**

**B, Translational trisomy 21 (Y9043, n=8/8).**

**C. Normal diploid (Y9044, n=7/7).**

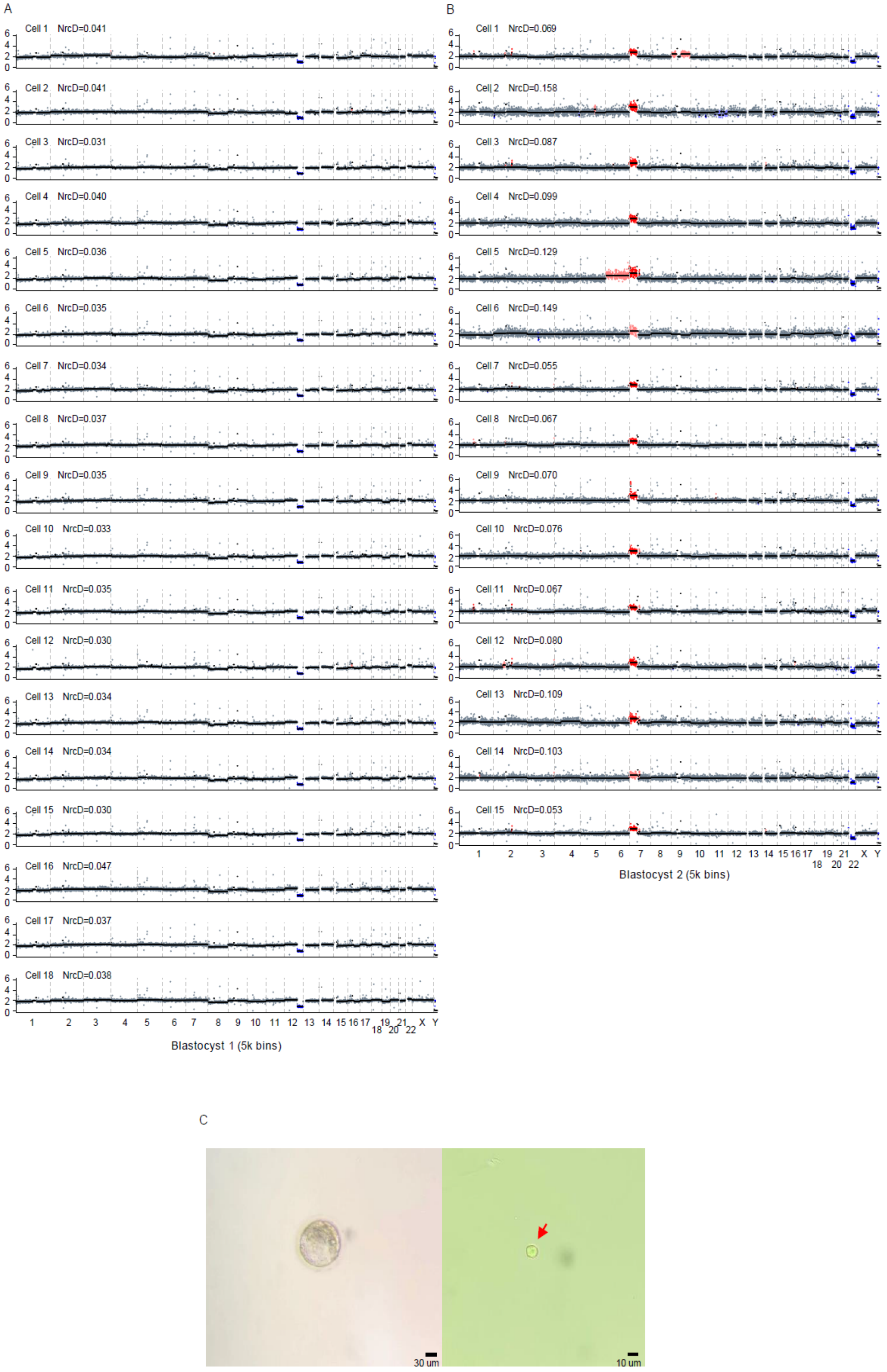
A
Cell 1 NrcD=0.041
Cell 2 NrcD=0.041
Cell 3 NrcD=0.031
Cell 4 NrcD=0.040
Cell 5 NrcD=0.036
Cell 6 NrcD=0.035
Cell 7 NrcD=0.034
Cell 8 NrcD=0.037
Cell 9 NrcD=0.035
Cell 10 NrcD=0.033
Cell 11 NrcD=0.035
Cell 12 NrcD=0.030
Cell 13 NrcD=0.034
Cell 14 NrcD=0.034
Cell 15 NrcD=0.030
Cell 16 NrcD=0.047
Cell 17 NrcD=0.037
Cell 18 NrcD=0.038
Blastocyst 1 (5k bins)
B
Cell 1 NrcD=0.069
Cell 2 NrcD=0.158
Cell 3 NrcD=0.087
Cell 4 NrcD=0.099
Cell 5 NrcD=0.129
Cell 6 NrcD=0.149
Cell 7 NrcD=0.055
Cell 8 NrcD=0.067
Cell 9 NrcD=0.070
Cell 10 NrcD=0.076
Cell 11 NrcD=0.067
Cell 12 NrcD=0.080
Cell 13 NrcD=0.109
Cell 14 NrcD=0.103
Cell 15 NrcD=0.053
Blastocyst 2 (5k bins)
C
30 um
10 um

**Extended Data Fig. 9. CNV patterns for single cells (blastomeres) from two preimplantation blastocysts by msCNVS.** The blastocysts were cultured on day 4, and discarded due to the poor quality (grade 4BC, retardated in development). The segment CNV is shown as follows. Blue: 1-copy or less; Purple: fuzzy deletion (value between 1-copy and 2-copy); Grey: 2-copy; Pink: fuzzy duplication (value between 2-copy and 3-copy); Red: 3-copy or more. Two blastocysts (A, B) show distinguished CNV patterns with chromosomal (Chr 21 in blastocyst 2) or segmental (sub-coromosomal) copy number duplication or deletion.

**A. msCNV calling for discarded blastocyst 1 (n=18).**

**B. msCNV calling for discarded blastocyst 2 (n=15).**

**C. Blastocyst 1 and a single cell dissociated from it.**

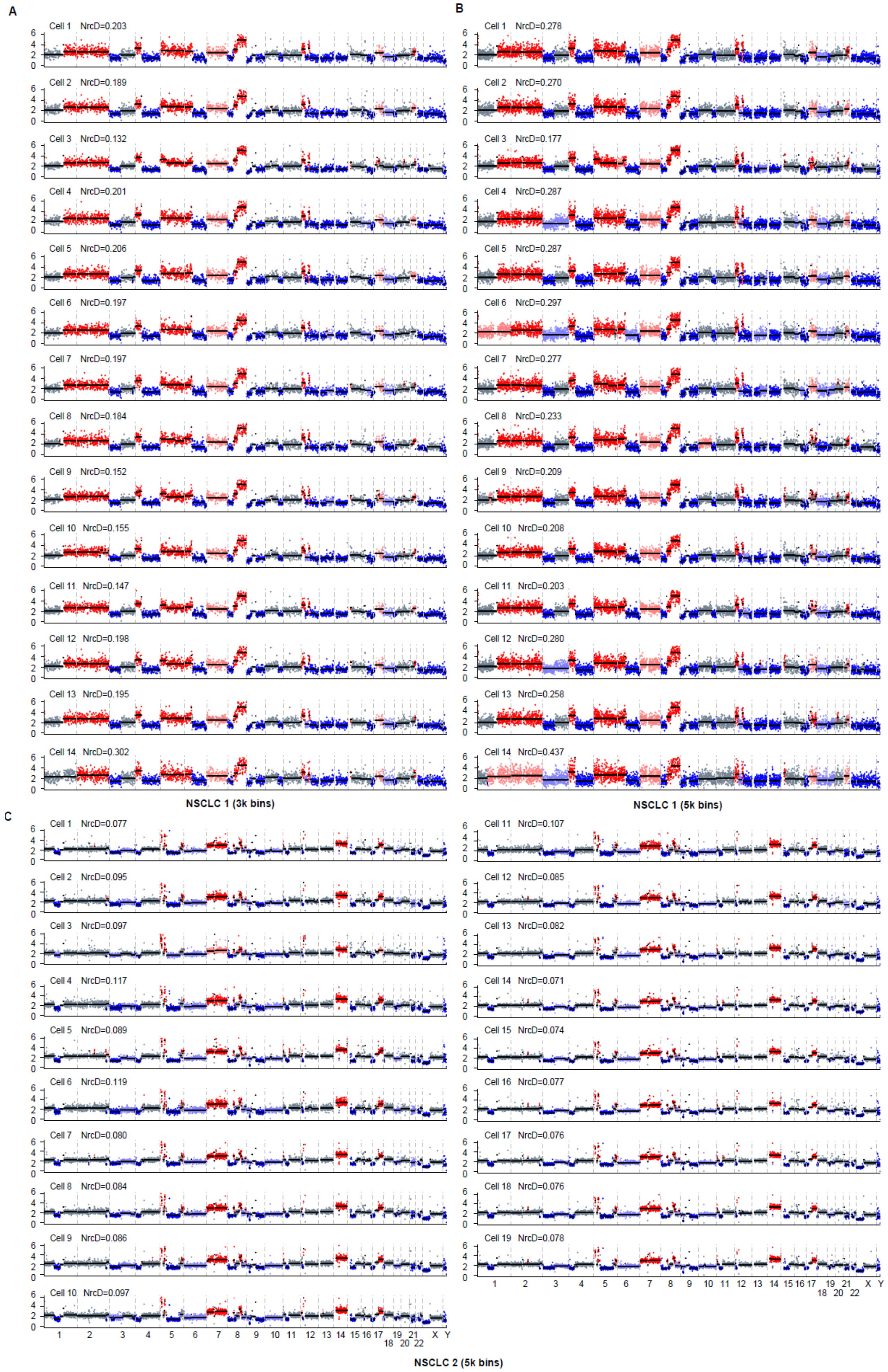
A
Cell 1 NrcD=0.203
Cell 2 NrcD=0.189
Cell 3 NrcD=0.132
Cell 4 NrcD=0.201
Cell 5 NrcD=0.206
Cell 6 NrcD=0.197
Cell 7 NrcD=0.197
Cell 8 NrcD=0.184
Cell 9 NrcD=0.152
Cell 10 NrcD=0.155
Cell 11 NrcD=0.147
Cell 12 NrcD=0.198
Cell 13 NrcD=0.195
Cell 14 NrcD=0.302
NSCLC 1 (3k bins)
B
Cell 1 NrcD=0.278
Cell 2 NrcD=0.270
Cell 3 NrcD=0.177
Cell 4 NrcD=0.287
Cell 5 NrcD=0.287
Cell 6 NrcD=0.297
Cell 7 NrcD=0.277
Cell 8 NrcD=0.233
Cell 9 NrcD=0.209
Cell 10 NrcD=0.208
Cell 11 NrcD=0.203
Cell 12 NrcD=0.280
Cell 13 NrcD=0.258
Cell 14 NrcD=0.437
NSCLC 1 (5k bins)
C
Cell 1 NrcD=0.077
Cell 2 NrcD=0.095
Cell 3 NrcD=0.097
Cell 4 NrcD=0.117
Cell 5 NrcD=0.089
Cell 6 NrcD=0.119
Cell 7 NrcD=0.080
Cell 8 NrcD=0.084
Cell 9 NrcD=0.086
Cell 10 NrcD=0.097
Cell 11 NrcD=0.107
Cell 12 NrcD=0.085
Cell 13 NrcD=0.082
Cell 14 NrcD=0.071
Cell 15 NrcD=0.074
Cell 16 NrcD=0.077
Cell 17 NrcD=0.076
Cell 18 NrcD=0.076
Cell 19 NrcD=0.078
1 2 3 4 5 6 7 8 9 10 11 12 13 14 15 16 17 19 21 X Y
18 20 22
NSCLC 2 (5k bins)

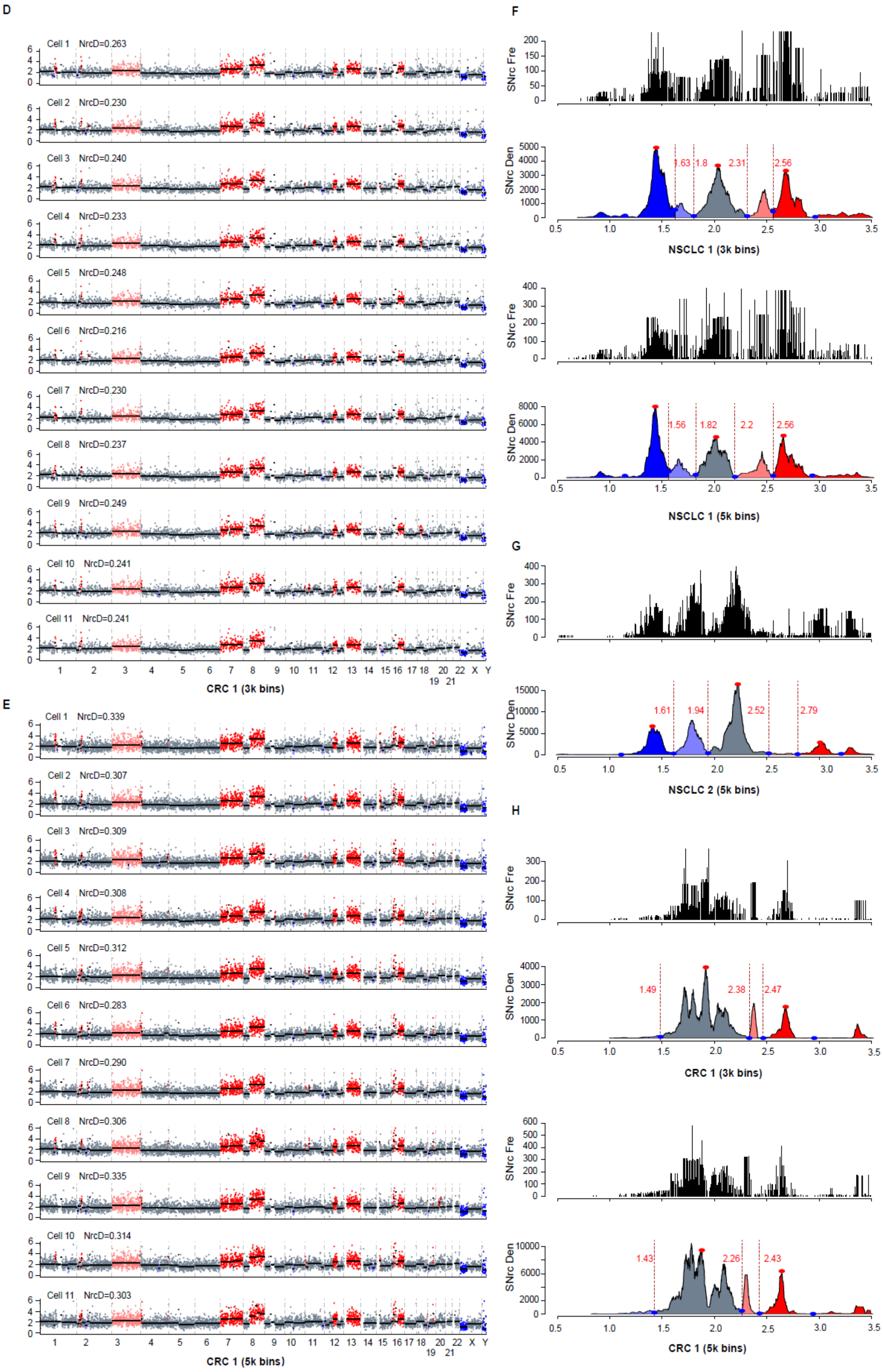
D
Cell 1 NrcD=0.263
Cell 2 NrcD=0.230
Cell 3 NrcD=0.240
Cell 4 NrcD=0.233
Cell 5 NrcD=0.248
Cell 6 NrcD=0.216
Cell 7 NrcD=0.230
Cell 8 NrcD=0.237
Cell 9 NrcD=0.249
Cell 10 NrcD=0.241
Cell 11 NrcD=0.241
CRC 1 (3k bins)
E
Cell 1 NrcD=0.339
Cell 2 NrcD=0.307
Cell 3 NrcD=0.309
Cell 4 NrcD=0.308
Cell 5 NrcD=0.312
Cell 6 NrcD=0.283
Cell 7 NrcD=0.290
Cell 8 NrcD=0.306
Cell 9 NrcD=0.335
Cell 10 NrcD=0.314
Cell 11 NrcD=0.303
CRC 1 (5k bins)
F
SNrc Fre
SNrc Den
1.63 1.8 2.31 2.56
NSCLC 1 (3k bins)
1.56 1.82 2.2 2.56
NSCLC 1 (5k bins)
G
1.61 1.94 2.52 2.79
NSCLC 2 (5k bins)
H
1.49 2.38 2.47
CRC 1 (3k bins)
1.43 2.26 2.43
CRC 1 (5k bins)

**Extended Data Fig. 10. CNV petterns of single circulated tumor cells (CTCs) from two patients of non-small cell lung cancer (NSCLC) and one patient of colorectal cancer (CRC).** CTCs were immune-captured and digital detected (see Materials and Methods). Generated by msCNVS, the CNV patterns among single CTCs within each patient are very similar, while they are distinguished between the two patients and two cancers. The segment CNV is shown as follows. Blue: 1-copy or less; Purple: fuzzy deletion (value between 1-copy and 2-copy); Grey: 2-copy; Pink: fuzzy duplication (value between 2-copy and 3-copy); Red: 3-copy or more.

**A/B. CNV pattern of CTCs from NSCLC 1 (n=14). A total of 14 CTCs are displayed. The NrcD is quallified at a resolution of 3k bins (B) for 13 CTCs, while most are non-qualified or at the edge with a resolution at 5k bins. It is noted that the Cell 14 is non-qualified with neither 3k bins and 5k bins.**

**C. CNV pattern of CTCs from NSCLC 2 (n=19).** A total of 19 CTCs are displayed. The NrcD is quallified at a resolution of 5k bins.

**D/E. CNV pattern of CTCs from CRC 1 (n=11/24). A total of 11 CTCs are displayed. The NrcD is quallified at a resolution of 3k bins (D) for 11 CTCs, while most are non-qualified or at the edge with a resolution at 5k bins (E).**

**F. SNrc FrequencyDistribution and SNrc Density distriubtion for the merged data of single CTCs from NSCLC 1 in 2 resolutions (3k bins versus 5k bins).**

**G. SNrc FrequencyDistribution and SNrc Density distriubtion for the merged data of single CTCs from NSCLC 2 in resolution of 3k bins.**

**H. SNrc FrequencyDistribution and SNrc Density distriubtion for the merged data of single CTCs from CRC 1 in 2 resolutions (3k bins versus 5k bins).**

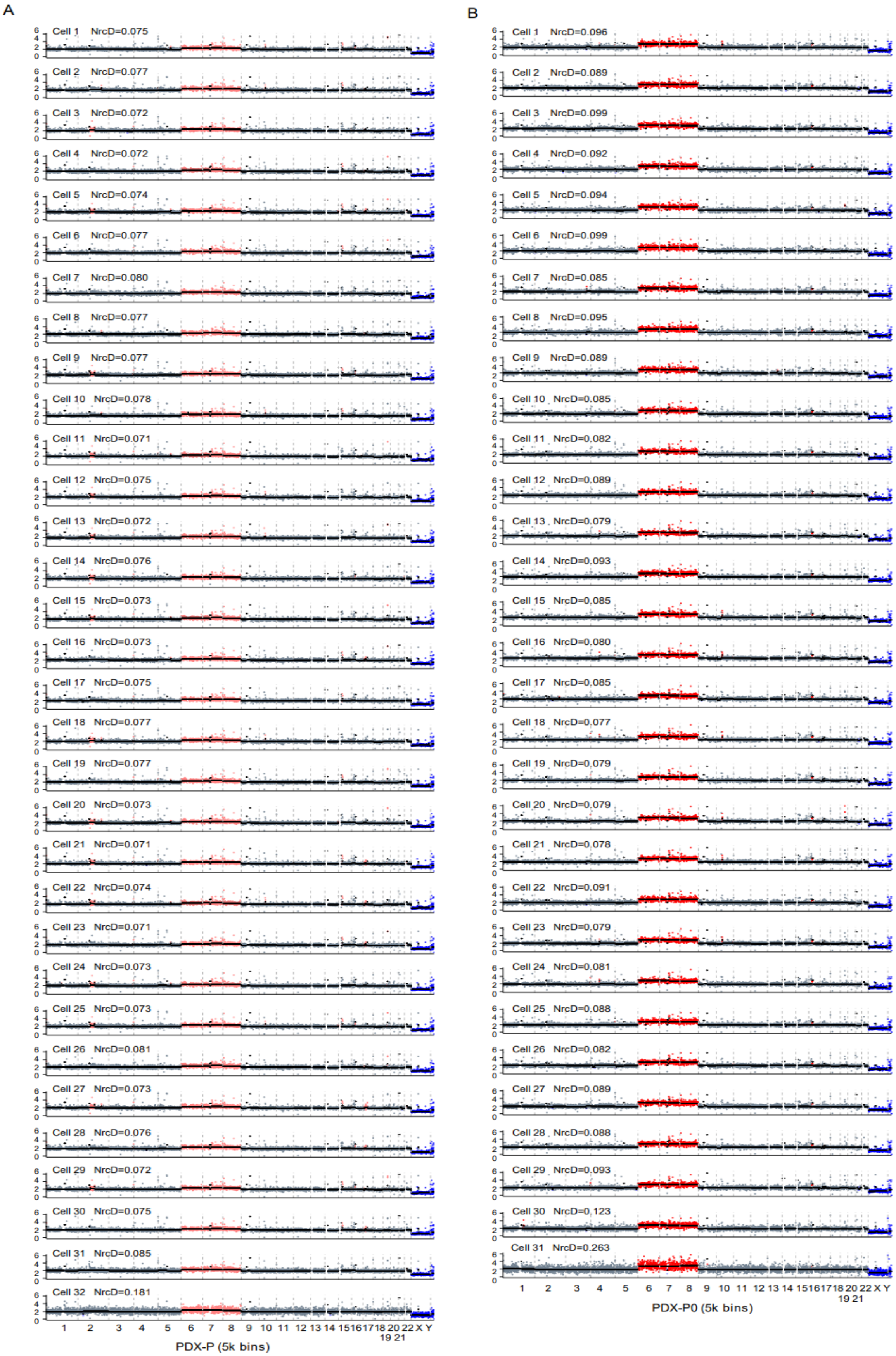
A
Cell 1 NrcD=0.075
Cell 2 NrcD=0.077
Cell 3 NrcD=0.072
Cell 4 NrcD=0.072
Cell 5 NrcD=0.074
Cell 6 NrcD=0.077
Cell 7 NrcD=0.080
Cell 8 NrcD=0.077
Cell 9 NrcD=0.077
Cell 10 NrcD=0.078
Cell 11 NrcD=0.071
Cell 12 NrcD=0.075
Cell 13 NrcD=0.072
Cell 14 NrcD=0.076
Cell 15 NrcD=0.073
Cell 16 NrcD=0.073
Cell 17 NrcD=0.075
Cell 18 NrcD=0.077
Cell 19 NrcD=0.077
Cell 20 NrcD=0.073
Cell 21 NrcD=0.071
Cell 22 NrcD=0.074
Cell 23 NrcD=0.071
Cell 24 NrcD=0.073
Cell 25 NrcD=0.073
Cell 26 NrcD=0.081
Cell 27 NrcD=0.073
Cell 28 NrcD=0.076
Cell 29 NrcD=0.072
Cell 30 NrcD=0.075
Cell 31 NrcD=0.085
Cell 32 NrcD=0.181
1 2 3 4 5 6 7 8 9 10 11 12 13 14 15 16 17 18 20 22 X Y
19 21
PDX-P (5k bins)
B
Cell 1 NrcD=0.096
Cell 2 NrcD=0.089
Cell 3 NrcD=0.099
Cell 4 NrcD=0.092
Cell 5 NrcD=0.094
Cell 6 NrcD=0.099
Cell 7 NrcD=0.085
Cell 8 NrcD=0.095
Cell 9 NrcD=0.089
Cell 10 NrcD=0.085
Cell 11 NrcD=0.082
Cell 12 NrcD=0.089
Cell 13 NrcD=0.079
Cell 14 NrcD=0.093
Cell 15 NrcD=0.085
Cell 16 NrcD=0.080
Cell 17 NrcD=0.085
Cell 18 NrcD=0.077
Cell 19 NrcD=0.079
Cell 20 NrcD=0.079
Cell 21 NrcD=0.078
Cell 22 NrcD=0.091
Cell 23 NrcD=0.079
Cell 24 NrcD=0.081
Cell 25 NrcD=0.088
Cell 26 NrcD=0.082
Cell 27 NrcD=0.089
Cell 28 NrcD=0.088
Cell 29 NrcD=0.093
Cell 30 NrcD=0.123
Cell 31 NrcD=0.263
1 2 3 4 5 6 7 8 9 10 11 12 13 14 15 16 17 18 20 22 X Y
19 21
PDX-P0 (5k bins)

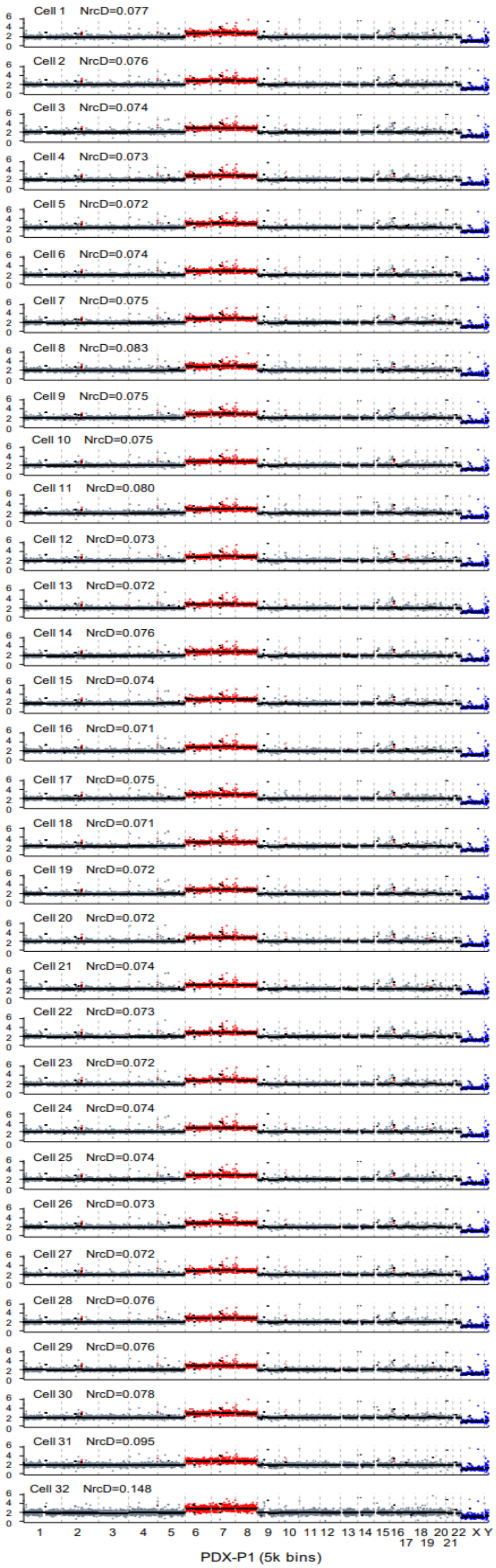
Cell 1 NrcD=0.077
Cell 2 NrcD=0.076
Cell 3 NrcD=0.074
Cell 4 NrcD=0.073
Cell 5 NrcD=0.072
Cell 6 NrcD=0.074
Cell 7 NrcD=0.075
Cell 8 NrcD=0.083
Cell 9 NrcD=0.075
Cell 10 NrcD=0.075
Cell 11 NrcD=0.080
Cell 12 NrcD=0.073
Cell 13 NrcD=0.072
Cell 14 NrcD=0.076
Cell 15 NrcD=0.074
Cell 16 NrcD=0.071
Cell 17 NrcD=0.075
Cell 18 NrcD=0.071
Cell 19 NrcD=0.072
Cell 20 NrcD=0.072
Cell 21 NrcD=0.074
Cell 22 NrcD=0.073
Cell 23 NrcD=0.072
Cell 24 NrcD=0.074
Cell 25 NrcD=0.074
Cell 26 NrcD=0.073
Cell 27 NrcD=0.072
Cell 28 NrcD=0.076
Cell 29 NrcD=0.076
Cell 30 NrcD=0.078
Cell 31 NrcD=0.095
Cell 32 NrcD=0.148
1 2 3 4 5 6 7 8 9 10 11 12 13 14 15 16 17 18 19 20 21 22 X Y
PDX-P1 (5k bins)

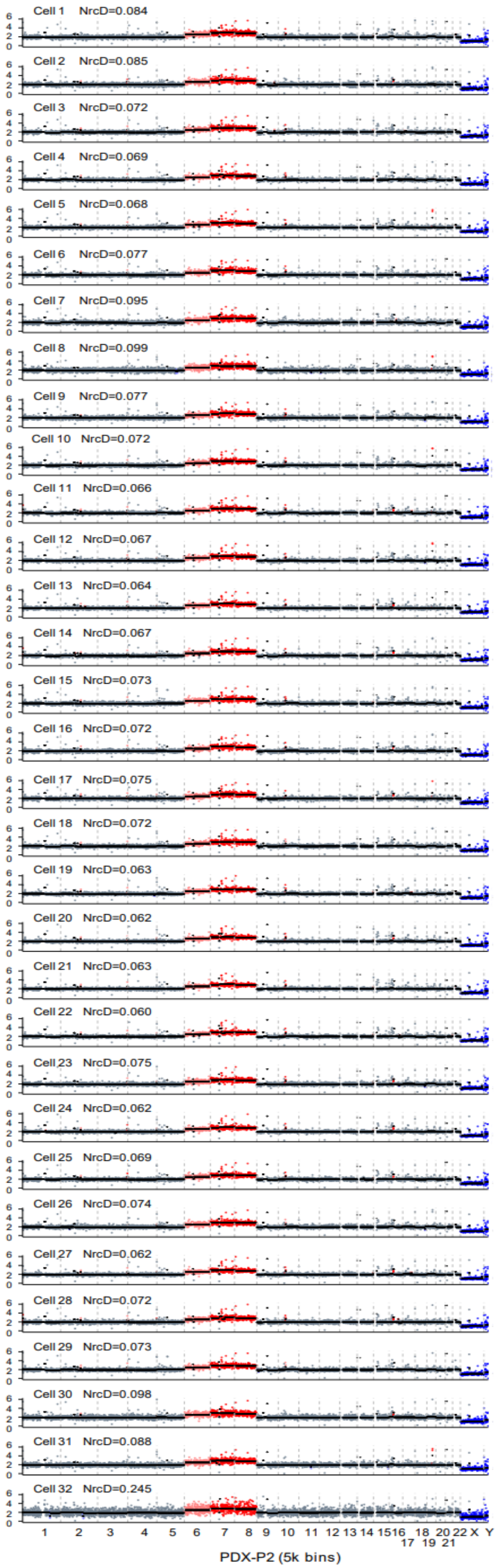
Cell 1 NrcD=0.084
Cell 2 NrcD=0.085
Cell 3 NrcD=0.072
Cell 4 NrcD=0.069
Cell 5 NrcD=0.068
Cell 6 NrcD=0.077
Cell 7 NrcD=0.095
Cell 8 NrcD=0.099
Cell 9 NrcD=0.077
Cell 10 NrcD=0.072
Cell 11 NrcD=0.066
Cell 12 NrcD=0.067
Cell 13 NrcD=0.064
Cell 14 NrcD=0.067
Cell 15 NrcD=0.073
Cell 16 NrcD=0.072
Cell 17 NrcD=0.075
Cell 18 NrcD=0.072
Cell 19 NrcD=0.063
Cell 20 NrcD=0.062
Cell 21 NrcD=0.063
Cell 22 NrcD=0.060
Cell 23 NrcD=0.075
Cell 24 NrcD=0.062
Cell 25 NrcD=0.069
Cell 26 NrcD=0.074
Cell 27 NrcD=0.062
Cell 28 NrcD=0.072
Cell 29 NrcD=0.073
Cell 30 NrcD=0.098
Cell 31 NrcD=0.088
Cell 32 NrcD=0.245
1 2 3 4 5 6 7 8 9 10 11 12 13 14 15 16 17 18 19 20 21 22 X Y
PDX-P2 (5k bins)

E

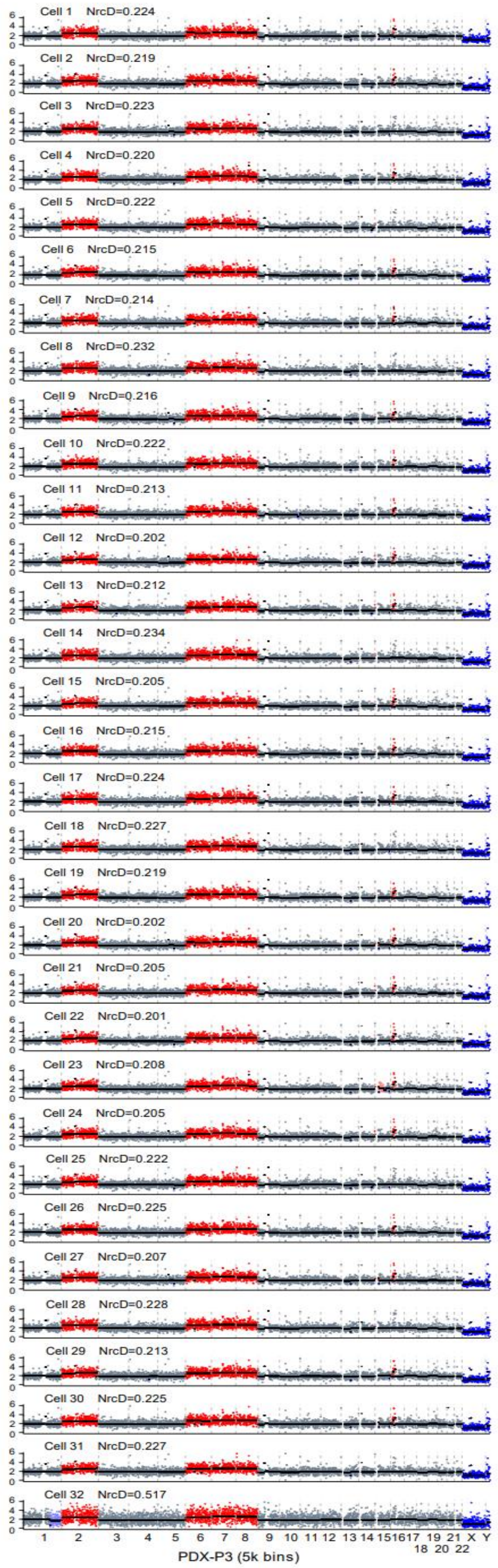
Cell 1 NrcD=0.224
Cell 2 NrcD=0.219
Cell 3 NrcD=0.223
Cell 4 NrcD=0.220
Cell 5 NrcD=0.222
Cell 6 NrcD=0.215
Cell 7 NrcD=0.214
Cell 8 NrcD=0.232
Cell 9 NrcD=0.216
Cell 10 NrcD=0.222
Cell 11 NrcD=0.213
Cell 12 NrcD=0.202
Cell 13 NrcD=0.212
Cell 14 NrcD=0.234
Cell 15 NrcD=0.205
Cell 16 NrcD=0.215
Cell 17 NrcD=0.224
Cell 18 NrcD=0.227
Cell 19 NrcD=0.219
Cell 20 NrcD=0.202
Cell 21 NrcD=0.205
Cell 22 NrcD=0.201
Cell 23 NrcD=0.208
Cell 24 NrcD=0.205
Cell 25 NrcD=0.222
Cell 26 NrcD=0.225
Cell 27 NrcD=0.207
Cell 28 NrcD=0.228
Cell 29 NrcD=0.213
Cell 30 NrcD=0.225
Cell 31 NrcD=0.227
Cell 32 NrcD=0.517
1 2 3 4 5 6 7 8 9 10 11 12 13 14 151617 19 21 X Y
18 20 22
PDX-P3 (5k bins)

**Extended Data Fig.11. Single nuclei of PDX derived from patient CRC 2.** The same CNV calling parameters apply to all samples except otherwise specified (F). A sum of 153 nuclei from the five generations of of PDX by msCNVS are displayed. The Chr 2 duplication appeared in the PDX bulk P3 (Fig. 5), which is outstanding over other generations, is also detected in the single cells of P3.

**A.Generation P (5k bins, n=32)**

**B.Generation P0 (5k bins, n=31)**

**C.Generation P1 (5k bins, n=32)**

**D.Generation P2 (5k bins, n=32)**

**E.Generation P3 (5k bins, n=32)**

**The CNV patterns of all generations are displayed at the resolution of 5k. The PDX-P3 may be paritally degrated and the NrcD values in PDX-P3 are relatively higher yet still within the threshold (0.25) except the cell# 32.**

### Tables S1 to S2:

**Table S1 Correlation between single cell and bulk in two resolution**

| single cell NO. | 5k bins | 10k bins | Resolution | SD | Median |
|---|---|---|---|---|---|
| 1 | 0.95145862 | 0.9206994 | 5k bins | 0.06459808 | 0.91044371 |
| 2 | 0.91331076 | 0.8653111 | 10k bins | 0.07444076 | 0.86871441 |
| 3 | 0.7986569 | 0.7414639 | | | |
| 4 | 0.83214592 | 0.8232244 | | | |
| 5 | 0.92358975 | 0.8841222 | | | |
| 6 | 0.90982527 | 0.7959288 | | | |
| 7 | 0.9414961 | 0.9247682 | | | |
| 8 | 0.92331386 | 0.8950279 | | | |
| 9 | 0.90136958 | 0.8708986 | | | |
| 10 | 0.92608515 | 0.9058782 | | | |
| 11 | 0.91953214 | 0.8881588 | | | |
| 12 | 0.92847953 | 0.9045319 | | | |
| 13 | 0.94203217 | 0.9171852 | | | |
| 14 | 0.93252415 | 0.906169 | | | |
| 15 | 0.85317899 | 0.7710052 | | | |
| 16 | 0.91704236 | 0.9026596 | | | |
| 17 | 0.83018064 | 0.7749054 | | | |
| 18 | 0.88749527 | 0.8594768 | | | |
| 19 | 0.8139621 | 0.7916249 | | | |
| 20 | 0.76189935 | 0.683064 | | | |
| 21 | 0.86677976 | 0.8448131 | | | |
| 22 | 0.90143305 | 0.8560891 | | | |
| 23 | 0.92017261 | 0.9055672 | | | |
| 24 | 0.90424068 | 0.8611842 | | | |
| 25 | 0.84503236 | 0.836606 | | | |
| 26 | 0.91266511 | 0.8881261 | | | |
| 27 | 0.91462407 | 0.877217 | | | |
| 28 | 0.85707634 | 0.7905247 | | | |
| 29 | 0.91308705 | 0.8958243 | | | |
| 30 | 0.92683082 | 0.8856055 | | | |
| 31 | 0.93001003 | 0.9072507 | | | |
| 32 | 0.88658958 | 0.8237918 | | | |
| 33 | 0.76036478 | 0.7036295 | | | |

| | | |
|---|---|---|
| 34 | 0.9004579 | 0.8472012 |
| 35 | 0.92611798 | 0.9066769 |
| 36 | 0.90184938 | 0.8821007 |
| 37 | 0.85505125 | 0.7633933 |
| 38 | 0.90357748 | 0.8852477 |
| 39 | 0.79081966 | 0.73431 |
| 40 | 0.8787539 | 0.8376915 |
| 41 | 0.8598997 | 0.7745212 |
| 42 | 0.81906754 | 0.7399279 |
| 43 | 0.93274703 | 0.8970749 |
| 44 | 0.91106215 | 0.7426369 |
| 45 | 0.91795078 | 0.8745635 |
| 46 | 0.8138059 | 0.7499779 |
| 47 | 0.89301394 | 0.810572 |
| 48 | 0.81901242 | 0.7922585 |
| 49 | 0.94233023 | 0.9247129 |
| 50 | 0.82415656 | 0.7753888 |
| 51 | 0.9295359 | 0.9164711 |
| 52 | 0.92813661 | 0.904786 |
| 53 | 0.94716155 | 0.9278983 |
| 54 | 0.90108993 | 0.8665302 |
| 55 | 0.80483367 | 0.7088798 |
| 56 | 0.57678509 | 0.5711738 |
| 57 | 0.92180436 | 0.8863816 |
| 58 | 0.79849057 | 0.7971862 |
| 59 | 0.91814294 | 0.8557279 |
| 60 | 0.9259657 | 0.8930131 |
| 61 | 0.92205747 | 0.910044 |
| 62 | 0.7225658 | 0.6737945 |
| 63 | 0.92761135 | 0.9177807 |
| 64 | 0.92466962 | 0.8826255 |
| 65 | 0.94644901 | 0.923125 |
| 66 | 0.8551012 | 0.8313145 |
| 67 | 0.9161354 | 0.8889341 |
| 68 | 0.92796352 | 0.8947822 |
| 69 | 0.93107201 | 0.8961608 |
| 70 | 0.81901798 | 0.7919355 |

### Table S2 Reads/windows in different resolution of K562 and HeLa by msCNVS

| | coverage(%) | mean depth | raw data | Total Reads | 100k | 200k | 400k | 600k | 800k | 1000k |
|---|---|---|---|---|---|---|---|---|---|---|
| **HeLa_bulk** | 31.659 | 0.629 | 36Gb | 13167567 | 396 | 799 | 1613 | 2424 | 3245 | 4080 |
| **K562_sc merge** | 38.833 | 0.897 | 76Gb | 24629032 | 751 | 1511 | 3044 | 4595 | 6115 | 7652 |
| **K562_cell1** | 4.599 | 0.068 | 3.3Gb | 1735035 | 53 | 107 | 215 | 323 | 433 | 541 |
| **K562_cell2** | 4.191 | 0.062 | 3.2Gb | 1577398 | 49 | 97 | 195 | 294 | 393 | 493 |
| **K562_cell3** | 4.528 | 0.062 | 1.4Gb | 1553539 | 47 | 95 | 190 | 287 | 383 | 480 |
| **K562_bulk** | 84.432 | 84.417 | 942G | 1229471315 | 42428 | 84857 | 169715 | 254571 | 339431 | 424285 |
